# Looking for the mechanism of arsenate respiration in an arsenate-dependent growing culture of *Fusibacter* sp. strain 3D3, independent of ArrAB

**DOI:** 10.1101/2022.06.08.495031

**Authors:** Acosta Grinok Mauricio, Susana Vázquez, Guiliani Nicolás, Sabrina Marín, Demergasso Cecilia

## Abstract

The literature has reported the isolation of arsenate-dependent growing (ADG) microorganisms which lack a canonical homolog for respiratory arsenate reductase, ArrAB. We recently isolated an ADG bacterium from arsenic-bearing environments in Northern Chile, *Fusibacter* sp. strain 3D3 (*Fas*) and studied the arsenic metabolism in this Gram-positive isolate. Features of *Fas* deduced from genome analysis and comparative analysis with other arsenic-reducing microorganisms revealed the lack of ArrAB coding genes and the occurrence of two *arsC* genes encoding for putative cytoplasmic arsenate reductases named ArsC-1 and ArsC-2. Interestingly, ArsC-1 and ArsC-2 belong to the thioredoxin-coupled family (because of the redox-active disulfide protein used as reductant), but they conferred differential AsV resistance to the *E. coli* WC3110 Δ*arsC* strain. PCR experiments confirmed the absence of *arrAB* genes and results obtained using uncouplers revealed that *Fas* growth is linked to the proton gradient. In addition, *Fas* harbors ferredoxin-NAD^+^ oxidoreductase (Rnf) coding genes. These are key molecular markers of a recently discovered flavin-based electron bifurcation mechanism involved in energy conservation, mainly in anaerobic metabolisms regulated by the cellular redox state and mostly associated with cytoplasmic enzyme complexes. At least three electron-bifurcating flavoenzyme complexes were evidenced in *Fas*, some of them shared in conserved genomic regions by other members of the *Fusibacter* genus. These physiological and genomic findings permit us to hypothesize the existence of an uncharacterized arsenate-dependent growth metabolism regulated by the cellular redox state in *Fusibacter* genus.

## Introduction

### The arsenic bioenergetic metabolisms

Until the 90’s, arsenic (As) was recognized only as a toxic compound for cells due to its role as a molecular analogous to phosphate, able to inhibit ATP synthesis, inactivate proteins and affect various intracellular processes. The new knowledge about the role of arsenic as a reactant in the bioenergetic microbial metabolisms has been summarized in 2014 by Amend et al. [1]. Anaerobic arsenic respiration using organic matter as electron donor [2, 3], energy conservation from bacterial arsenite oxidation [4, 5] and the use of arsenite as a fuel for anoxygenic photosynthesis complement the description of the microbially-catalyzed arsenic metabolisms [6, 7].

The pathways involving As metabolizing enzymes contribute to the generation of chemiosmotic potential by coupling exergonic electron transfer to proton translocation with As playing the role of oxidizing (arsenate reductase, Arr) or reducing (arsenite oxidase Aio, or the alternative arsenite oxidase Arx, a variant of Arr but working in reverse) substrates [1]. Arr operates by funneling reducing equivalents from organic matter to the terminal acceptor AsV in an anaerobic respiration involving the quinol pool. According to the possible metabolic pathways, Aio transfers electrons from AsIII towards O_2_, NO^-^, chlorate or the photosynthetic reaction center through a chain of electron carriers: Cyt and Cox, Fdh-Nar-Nir-Nor-NosZ, soluble cytochrome and cytochrome-chlorate oxide-reductase (Clr) or the membrane-bound auracyanin, respectively. In 2013, van Lis et al. (2013) stipulated that all the Arr-harboring strains oxidize the liposoluble menaquinone (MK) pool via an AsV reduction process whereas the Arx-harboring strains reduce the ubiquinone (UQ) pool via an AsIII oxidation process [8]. MK and UQ biosynthetic pathways were clearly identified in Arr and Arx-harboring strains, respectively. Besides that, it has been reported that some AsV reducing microorganisms, such as *Shewanella* sp. strain HN-41, *Desulfomicrobium* sp. strain Ben-RB, *Citrobacter* sp. strain TSA-1, *Fusibacter* sp. strain 3D3 and *Pyrobaculum aerophilum* strain IM2 do not contain *arrAB* genes [1, 9–14], indicating that there should be at least one alternative and uncharacterized pathway for arsenic reduction.

### The arsenic resistance mechanism

Many bacteria can detoxify As by the plasmid or chromosomal encoded Ars system, which is widespread in nature and has been extensively studied [15, 16]. The arsenical resistance (*ars*) operon includes up to five genes, among which *arsR*, a transcriptional regulator encoding gene,)and the *arsC* gene are almost always present [17]. The key enzyme is ArsC, a cytoplasmic arsenate reductase reducing AsV to AsIII, which is then extruded out of the cell by the pump coded by the *arsB* gene. Three different ArsC prokaryotic families have been defined based on their protein structures, reduction mechanisms and location of the catalytic cysteine residues [18]: i) the glutathione (GSH)/glutaredoxin (Grx)-coupled class (plasmid R773 from Gram-negative bacteria *Escherichia coli*) [19]; ii) the thioredoxin (Trx)/thioredoxin reductase (TrxR)-dependent class (plasmid pl258 from Gram-positive bacteria *Staphylococcus aureus*) [20]; iii) the mycothiol (MSH)/mycoredoxin (Mrx)-dependent class (chromosome of Gram-positive *Actinobacteria* spp) where MSH is the major thiol [21]. Kinetics data of arsenate reduction have shown a higher catalytic efficiency in Trx-than in Grx-linked arsenate reductases. In some cases, the efflux of AsIII can also be coupled to the electrochemical proton gradient, where chemical energy in the form of ATP is used to pump AsIII with the help of the ATPase ArsA [22–25].

Both, the number and type (Trx or Grx clade) of *arsC* genes present in the genomes of prokaryotic organisms have been shown to impact their As resistance levels [18, 26–28]. The Trx reducing system has been reported to be the most efficient system exploited by arsenate reductases [18]. Besides, arsenate reductases with the same structural fold but depending on two different thiol-disulfide relay mechanisms (Trx and GSH) have also been observed in a single microorganism, *Corynebacterium glutamicum* ATCC 13032 [18]. In that case, a different role has been proposed for both enzymes, representatives of different ArsC prokaryotic families: the Trx-dependent would reduce arsenate to regulate the gene expression of the other one, that is involved in the resistance against As [18]. A predominance of Trx-linked ArsC have been found in low G+C Gram-positive bacteria [29], which is the predominant group of bacteria in the As-impacted environments of Northern Chile [30–32].

### Another energy conservation mode: “A new era for electron bifurcation”

In 2008, a third type of energy conservation mode [33] in addition to substrate-level phosphorylation (SLP) and electron transport phosphorylation (ETP) has been discovered, almost exclusively associated with anaerobic metabolism. This type of energy conservation is based on two main components: i) the flavin-based electron bifurcation (FBEB), where ferredoxins and flavodoxins act as low-potential terminal acceptors, and ii) the ETP with protons (ferredoxin-proton reductase, Ech) or NAD^+^ (ferredoxin-NAD^+^ reductase, Rnf) as electron acceptors, where ferredoxins and flavodoxins re-oxidation drive electrochemical H^+^ and Na^+^ pumps. This energy conservation system has allowed closing gaps between the free energy change and the number of ATP molecules synthesized (ATP/ΔG) in the energy metabolism of some anaerobes [34, 35].

In many acetogens, acetoclastic methanogens, sulfate reducers and other strict anaerobes [36–39], the Rnf complex catalyzes the reversible oxidation of reduced ferredoxin with NAD^+^, coupling this exergonic reaction with the build-up of an electrochemical proton or sodium ion potential [34]. In different species, it has been demonstrated that the Rnf complex can be associated to the generation of either a Na^*+*^ gradient [40] or of a H^+^ gradient [41]. The stoichiometry is most likely one sodium ion or proton translocated per electron [39]. Consistent with the finding of a Na^+^-dependent Rnf complex, a conserved Na^+^-binding motif in the ATP synthase has been reported [42] and when a H^+^-dependent Rnf complex was found, a conserved H^+^-binding motif in the ATP synthase was reported instead [43]. The Rnf complex was first discovered in *Rhodobacter capsulatus* [44] and has a high sequence similarity with the Na^+^-translocating NADH:quinone-oxidoreductase (Nqr) [44, 45]. Some genomic, transcriptomic and proteomic reports have described the subunits conforming the RnfAG-complex, which are variable in number and organization depending on the species, as well as their main role in the generation of membrane electrochemical gradients and, therefore, in energy conservation [36, 46, 47]. Briefly, each subunit of the Rnf complex has a specific function related to cytosolic ferredoxin oxidation (RnfB), proton/sodium membrane translocation (RnfGD) or NAD^+^ cytosolic reduction (RnfC) as has been described in several microorganisms such as *Acetobacterium woodii, Escherichia coli, Clostridium tetani, Methanosarcina acetivorans*, and *Desulfovibrio aleskensis*, among others [48, 49].

In the same way, up to twelve multienzyme complexes involved in the electron bifurcation (referred to as electron confurcation when operating in reverse) mechanism and associated to energy conservation have been reported according to the 2018 and 2019 reviews on the subject [39, 50–53]. All known electron bifurcating enzymes contain at least one flavin cofactor (FAD or FMN), and it is why the novel mechanism was defined as a FBEB mechanism [53]. Interestingly, the distribution studies of the identified FBEB enzymes have shown that they are predominantly present among members of the *Firmicutes* and contribute to diverse metabolic pathways [52, 53]. A key role in balancing the ratio of oxidized to reduced NAD(H) and ferredoxin (Fd) pools has been proposed for the FBEB mechanism and its presence in arsenate reducers has also been identified (e.g., *Alkaliphilus oremlandii* OhILAs) [52].

Functional confirmation of energy conservation by the Rnf-mediated electron transport chain has been recently reported [41]. The *rnfAB*-mutants of the anaerobe *Clostridium ljungdahlii* were unable to grow on H_2_/CO_2_, demonstrating the important role of the Rnf complex in pumping H^+^ out of the cell membrane for energy conservation during autotrophic growth. ATP synthesis was also significantly reduced in the *rnfAB*-mutants during heterotrophic growth on fructose. Moreover, in the acetogenic *Acetobacterium woodi*, the sequence of events reported to be compatible with the caffeate reduction coupled to ATP synthesis [40] is: caffeate reduction ⟶ generation of a transmembrane Na^+^ gradient ⟶ generation of ATP by the Na^+^ F0F1 ATP synthase. The role of the Rnf complex in the Na^+^-dependent electron transfer reaction from reduced ferredoxin to NAD^+^ and vice versa was confirmed at the functional level in *A. woodii* by means of the protein and enzymatic activity assays and by genetic evidence [46, 54]. Another functional confirmation was reported in the sulfate reducer *Desulfovibrio alaskensis*, where *rnfA* and *rnfD* null-mutants were unable to grow on H_2_, formate and ethanol [55], as reported in other studies [41, 56]. Moreover, an increased expression level of *rnf* genes was observed in *D. alaskensis* growing with H_2_ and sulfate compared to lactate and sulfate. Some authors have inferred that *D. alaskensis* likely relies extensively on ferredoxin oxidation by the Rnf complex to produce a H^+^ gradient during growth on substrates that do not yield ATP by SLP [55, 56]. However, for those substrates that do yield ATP by SLP such as malate, fumarate, pyruvate and lactate, a decreased growth rate and/or yield was also observed in most cases for the *rnf* mutants [55]. Finally, it has been reported that the lactate dehydrogenase-electron transferring flavoprotein complex, the ferredoxin and the Rnf complex are key components in the lactate metabolism of *A. woodii*, a strictly anaerobic bacteria [57].

Altogether, these data pointed out that the Rnf complex is pivotal for anaerobic growth in microorganisms without cytochromes, quinones or other membrane-soluble electron carriers. However, it has also been proposed that metals like molybdenum, transition metal ions with three readily accessible oxidation states under *in vivo* conditions, could also be the site of electron bifurcation, of which the molybdenum in the arsenite oxidases could be one example [50].

### Arsenate respiration independent of ArrAB

The absence of a homolog for the respiratory arsenate reductase gene, a*rrAB*, has been reported for other strains. In one of them, *Pyrobaculum aerophilum*, a high expression of a gene cluster encoding for a molybdopterin oxidoreductase (PAE1265) with a molybdopterin-binding subunit was observed in cultures induced with arsenate. The analysis of the predicted product of PAE1265 showed the occurrence of an active site domain conserved in bacterial tetrathionate reductases and arsenate reductases [12, 58], leading to the hypothesis that tetrathionate reductases may represent a novel type of respiratory arsenate reductases. Based on this hypothesis, Blum and collaborators [10] performed a transcriptomic analysis focused on the tetrathionate reductase (*ttrA*), *arsC*, and 16S rRNA genes from *Citrobacter* TSA grown on arsenate. They reported that only *arsC* mRNA was strongly expressed, while there was a little detectable upregulation for *ttrA*, proposed to be the mean to achieve dissimilatory arsenate reduction. After those experimental results, they hypothesized that “*it is possible that there is an electron flow linkage to the detoxifying ArsC protein serving in a unique respiratory capacity and presumably located in the membrane region rather than the cytoplasm*”. Interestingly, similar gaps in the energy metabolism of anaerobes [34, 35] were closed by the characterization of the energy conservation system depending on a FBEB ferredoxin reduction and on a proton/sodium translocating ferredoxin oxidation [38]. This is considered as a third type of energy conservation mode [34] in addition to SLP and ETP.

### *Fusibacter* sp. strain 3D3 as a case study

Our research group has confirmed the occurrence of an active As biogeochemical cycle in Salar de Ascotán from metagenomic analysis [32]. Prokaryotic populations compatible with microorganisms able to transform As for energy conservation to produce H_2_, H_2_S and acetic acid (potential electron sources for As reduction) and tolerate high levels of As by means of specific stress response are involved in this cycle. Furthermore, the characterization of enrichment cultures confirmed their ability to metabolize As [32], some of its components being members of genera that, like *Fusibacter*, had not been previously reported as As metabolizing microorganisms [13, 59]. There is no report of arsenic resistance in other isolates of the *Fusibacter* genus, however, microorganisms closely related to this genus have been reported to occur in contaminated groundwater in Bangladesh [60]. The genome sequence of *Fas* [13] had suggested the presence of an *arsC* gene.

Then, the aim of this work was to describe the role of the cytoplasmic arsenate reductase (ArsC) and the membrane-associated ion-translocating complex (Rnf) in the energy metabolism of *Fusibacter* sp. strain 3D3, as a representative of arsenate reducing microorganisms independent of ArrAB.

## Materials and methods

### Bacterial strains

*Fusibacter* sp. strain 3D3 was isolated at the Centro de Biotecnología, Universidad Católica del Norte, Antofagasta, Chile, from samples collected in the hypersaline sediments of the Salar de Ascotán in Northern Chile and deposited in the American Type Culture Collection as *Fusibacter ascotence* ATCC BAA-2418 (hereinafter referred to as *Fas*). The necessary tests and deposits to describe the isolate as a new species are running, and “*Fusibacter ascotence*” is the proposed name. The *Fas* genome assembly [13] is available on NCBI (RefSeq GCF_001748365.1, GenBank GCA_001748365.1). The *E. coli* WC3110 Δ*arsC* strain was generously given by Dr. Barry P. Rosen.

### Culture characterization

All growth experiments were performed in duplicate in serum bottles containing 20 mL liquid Newman-modified minimal medium with lactate (10 mM), sulfate (20 mM), arsenate (2 m M), yeast extract (0.1%), NaCl (10 g L^-1^), and cysteine (1 mM), inoculated with 1×10^6^ cells mL^-1^ from a fresh culture and incubated at 30 °C in an anaerobic chamber under N_2_:CO_2_:H_2_ gas atmosphere (80:15:5, v/v) for 5 to 10 days in the dark, unless otherwise stated. An abiotic control was carried out in sterile medium without inoculum. Cell growth was monitored by microscope cell counting using a Neubauer improved chamber (0.01 mm x 0.0025 mm^2^, Marienfeld). To test for growth in the presence of oxygen, aerobic culture s were performed in shaking flasks incubated at 100 rpm in a rotatory shaker. The range of temperature fo r growth was tested between 15 °C and 37 °C, and the range of pH between 4 and 9. To assess the ferment ative metabolism, *Fas* was grown with alternative substrates: lactate, acetate, citrate, glucose, galactose, glycine, or tryptone (10 mM). Sodium thiosulfate (10 mM), sodium sulfate (0 to10 mM), elemental sulfur (1%), yeast extract (0.2 %) or cysteine (1 mM) were added to culture media to determine its ability to obtain energy from sulfate reduction and to use different sulfur sources. To test for AsV resistance and the opti mal concentration of As for energy metabolism, a range of concentrations between 0 and 16 mM was assa yed. To find out if the growth of *Fas* on AsV as electron acceptor was linked to oxidative phosphorylation and formation of proton or sodium gradients, growth was also evaluated with the addition of the protonophore 3,3’, 4’,5-tetrachlorosalicylanide (TCS, 20 μM) or the sodium-specific ionophore N,N,N’,N’-tetracyclohexyl-1,2-phenylenedioxydiacetamide (ETH2120, 20 μM) [41, 61].

### Analytical methods

To evaluate the arsenic and sulfate reducing activity, As concentrations in the culture medium from bacterial cultures were measured, after filtering through 0.02 μm pore size, using Hydride Generation Atomic Absorption Spectroscopy (HG-AAS) and the As speciation, AsIII and AsV, was analyzed using a Chromatography PSA 10.055 Millennium Excalibur. Lactate, acetate, and sulfate were quantified by ion chromatography (Dionex) with an IONPAC AS11-HC analytical column (4×250).

### Trx and TrxR enzymatic assays

To evaluate the participation of the Trx system in the early (30 min) and late (8 h) response to As exposure, the TrxR and Trx activities were measured at 30 °C using whole cell extracts as described previously [62], and a control without As exposure was also included in the experiment. TrxR was assayed for reductive activity toward 5,5-dithio-bis-(2-nitrobenzoic acid) (DTNB) with NADPH to form 5-thio-2-nitrobenzoic acid (TNB), producing a strong yellow color that was measured at 412 nm [63]. Total Trx activity was determined by the insulin precipitation assay [64]. The standard assay mixture contained 0.1 M potassium phosphate (pH 7.0), 1 mM EDTA, and 0.13 mM bovine insulin in the absence or in presence of the cellular extract, the reaction was started upon the addition of 1 mM DTT and the increase of the absorbance at 650 nm was monitored.

### DNA purification

Bacterial DNA was extracted and purified using the High Pure PCR Template Preparation kit (Roche, cat. n° 11796828001) according to the manufacturer’s protocol. PCR products were purified using the QIAquick PCR Purification kit (QIAGEN, cat. n° 28104), and digested DNA products were extracted from the agarose gel using the QIAquick Gel Extraction kit (QIAGEN, cat. n° 28704), according to the manufacturer’s instructions.

### PCR conditions

The presence of an arrA gene in the Fas genome was assessed by a PCR assay performed using the universal arrAf and arrAr primers to target an arrA internal ∼160-200 bp DNA fragment, as previously described [65]. Genomic DNA from Shewanella sp. strain ANA-3 was used as a positive control. To clone the Fas arsC-1 and arsC-2 genes, specific primers were designed based on both nucleotide sequences obtained from Fas genome and modified to include XhoI and HindIII restriction sites (Table S1). The PCR assays were performed with the Phusion High-Fidelity DNA Polymerase (Thermo Scientific, F-350S) according to the manufacturer’s instructions. The following PCR conditions were used: initial denaturation at 95 °C for 30 s, followed by 30 cycles of 98 °C for 10 s, 56.1 °C (arsC-1Fas) or 60.0 °C (arsC-2Fas) for 25 s, and 56.1 °C (arsC-1Fas) or 60.0 °C (arsC-2Fas) for 1 min, and a final elongation at 56.1 or 60 °C for 3 min. Then PCR products were run by electrophoresis and purified from the 1.5% agarose gel. Screening for recombinant colonies was made by PCR as previously described [66] using the PCR conditions described above and the GoTag kit (Promega, M3001).

### Cloning, heterologous expression of *arsC* genes and evaluation of AsV resistance

To confirm if *Fas* ArsC confer resistance to AsV, the *arsC*-1_*Fas*_ and *arsC-*2_*Fas*_ genes were amplified as described above. The purified PCR products were subjected to A-tailing with Taq DNA polymerase and to ligation into a T-vector (pGEM®-T Easy Vector, Promega, cat. n° A1360). The ligation product was used to transform *E. coli* JM109 (Competent Cells, Promega, cat. n° L2005) and recombinant clones were checked by sequencing. Plasmids with the *arsC*-1_*Fas*_ and *arsC-*2_*Fas*_ correct sequences were purified by miniprep (Wizard® Plus SV Minipreps, Promega, cat. n° A1360) and double digested with *Xho*I and *Hind*III (Thermo Scientific^™^, cat. n° ER0691 and ER0501, respectively) to release the inserts, which were subsequently purified and ligated upstream the His-tag into the expression vector pTrcHis2 (Invitrogen™, cat. n° V36520). The recombinant vectors were transformed into the Δ*arsC E. coli* WC3110 strain. Positive clones were cultured in LB medium with ampicillin 50 μg/mL for 12 h at 37 °C. The expression of both *arsC* genes was induced with IPTG 1 mM and the conferred ability to growth in the presence of 0, 0.5, 1, 1.5, 2.5 and 5 mM AsV was monitored by OD_600_ compared to a clone of *E. coli* WC3110 transformed with the pTrc-*lacZ* used as control. The specific growth rate (µ) was calculated with the equation: μ=(lnX-lnX_0_)/(t-t_0_), where X and X_0_ represent the OD_600_, and t and t_0_ the time. The doubling time (t_d_) was determined using the equation: t_d_=ln2/μ.

### Bioinformatic analysis

The genomes of *Fusibacter* sp. strain 3D3 (BDHH00000000.1), *F. ferrireducens* strain Q10-2^T^ (JADKNH000000000.1), *F. paucivorans* strain SEBR 4211^T^ (JAHBCL000000000.1), *F. tunisiensis* strain BELH1^T^ (JAFBDT000000000.1), *Fusibacter* sp. strain A1 (JABKBY000000000.1) were obtained from the NCBI database and annotated on the RAST platform using default settings [67].

The genome sequence of *Fas* [13] was screened to search for genes encoding components and regulators of As redox and transport system, as well as those associated with thiol redox systems and energy metabolisms. Curation of genes of interest was performed by reciprocal analysis against each other and the sequences available in the public databases to establish similarities and differences regarding gene identity, structure and function, gene context and control signals [68–72]. The sequence collections available on the NCBI [73] and the Comprehensive Microbial Resource of the J. Craig Venter Institute (Rockville, MD, USA) [74] websites facilitated comparative genomic studies with other organisms of interest whose genome sequencing had already been completed.

For the verification of the EtfB (electron transfer flavoprotein subunit beta) domain conservation across *Fusibacter*, previously known proteins from *Geobacter metallireducens* GS-15, *Thermotoga maritima* MSB8, *Clostridium ljungdahlii* PETC, *Rhodopseudomonas palustris* BisA53, *Acetobacterium woodii* WB1, *Clostridium kluyveri* DSM 555, *Acidaminococcus fermentans* VR4, and *Megasphaera elsdenii* T81 were used [75]. The protein sequences were aligned by MUSCLE [76] and visualized by CLC Genomic Workbench 8.5.1 (Qiagen). Subsequently, using BLAST on the RAST platform, EtfB-type proteins were searched in *Fusibacter* genomes, and the results were verified by BLASTp in the NCBI database. A similar procedure was applied to the products of the remaining genes.

## Results

### Microbial growth

*Fas* grew optimally at temperatures ranging from 20 to 37 °C, with the lower duplication time (t_d_) observed at 30 °C (Fig. 1A) and under neutral (6-8) pH conditions (Fig. 1B). *Fas* grew in synthetic medium containing up to 50 g L^-1^ of NaCl, and the optimum was 10 g L^-1^ [13]. No growth differences were detected at increasing SO_4-2_ concentrations (Fig. 1C). The highest specific growth rate (μ) was observed at 2 mM of AsV and pH 7 in anaerobiosis, but it grew in the range of 0.5 to up to 16 mM of AsV (Fig.1D). The lowest concentration of AsV that completely prevented growth (MIC) of *Fas* was 24 mM. Liquid cultures reached total bacterial numbers between 3.5×107 and 6.5×107 cells mL-1 at the stationary phase (data not shown).

**Figure 1.**
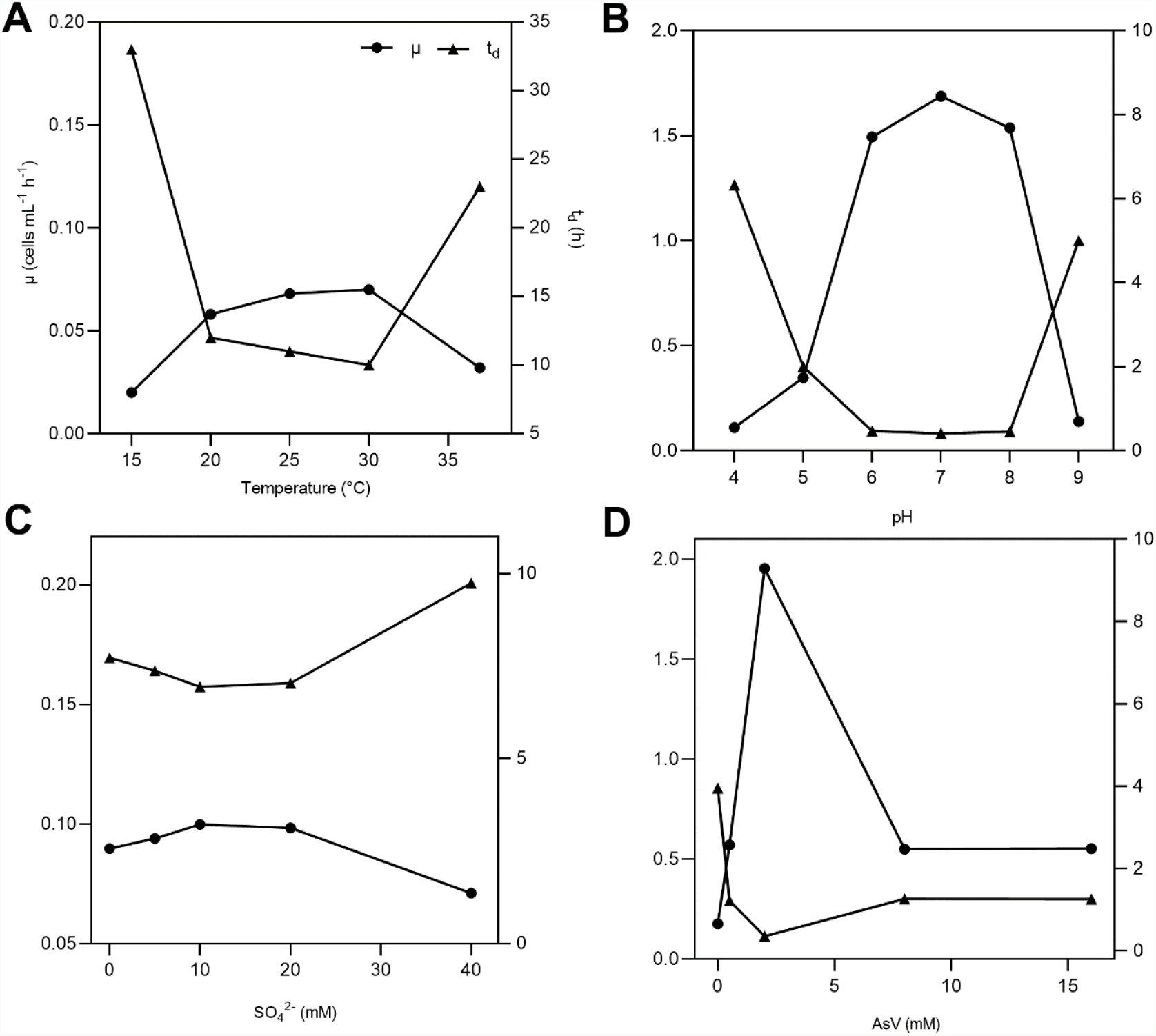
Growth of *Fas* in arsenate-containing medium. Duplication time (t_d_) and specific growth rate (μ) of *Fas* growth on increasing temperatures (A), pH (B), concentrations of SO_4_^2-^ (C) and AsV (D). Optimum temperature (30°C), pH (7), and concentrations of As (2 mM) and sulfate (20 mM) were used in the experiments except when different values of the corresponding variables were analyzed.

Not a significant change was noticed in the doubling time in minimal medium plus arsenate (2 mM) by the addition of sulfate (0 to 20 mM) (Fig. 1C) [13]. Beside, in minimal medium plus arsenate (2 mM) and sulfate (20 mM) as electron acceptors *Fas* grew with a doubling time of 0.35 h, which increased to 4 h when As was not supplemented (Fig. 1D).

### Features of *Fusibacter* sp. strain 3D3 metabolism

#### Substrates and products

The strain was not able to grow in aerobiosis but grew in anaerobiosis on lactate in the presence of sulfate and arsenate as electron acceptors. Defined as a heterotrophic strain, *Fas* could use lactate (Fig. 2), glucose and tryptone and required yeast extract to grow (Table 1). Growth without the addition of electron acceptors was successful up to the second subculture (data not shown) as it was in the previously reported culture medium for *Fusibacter* [77]. Besides, the highest AsV to AsIII reduction ratio was evidenced between 72 and 96 hours (Fig. S1).

**Table 1.**
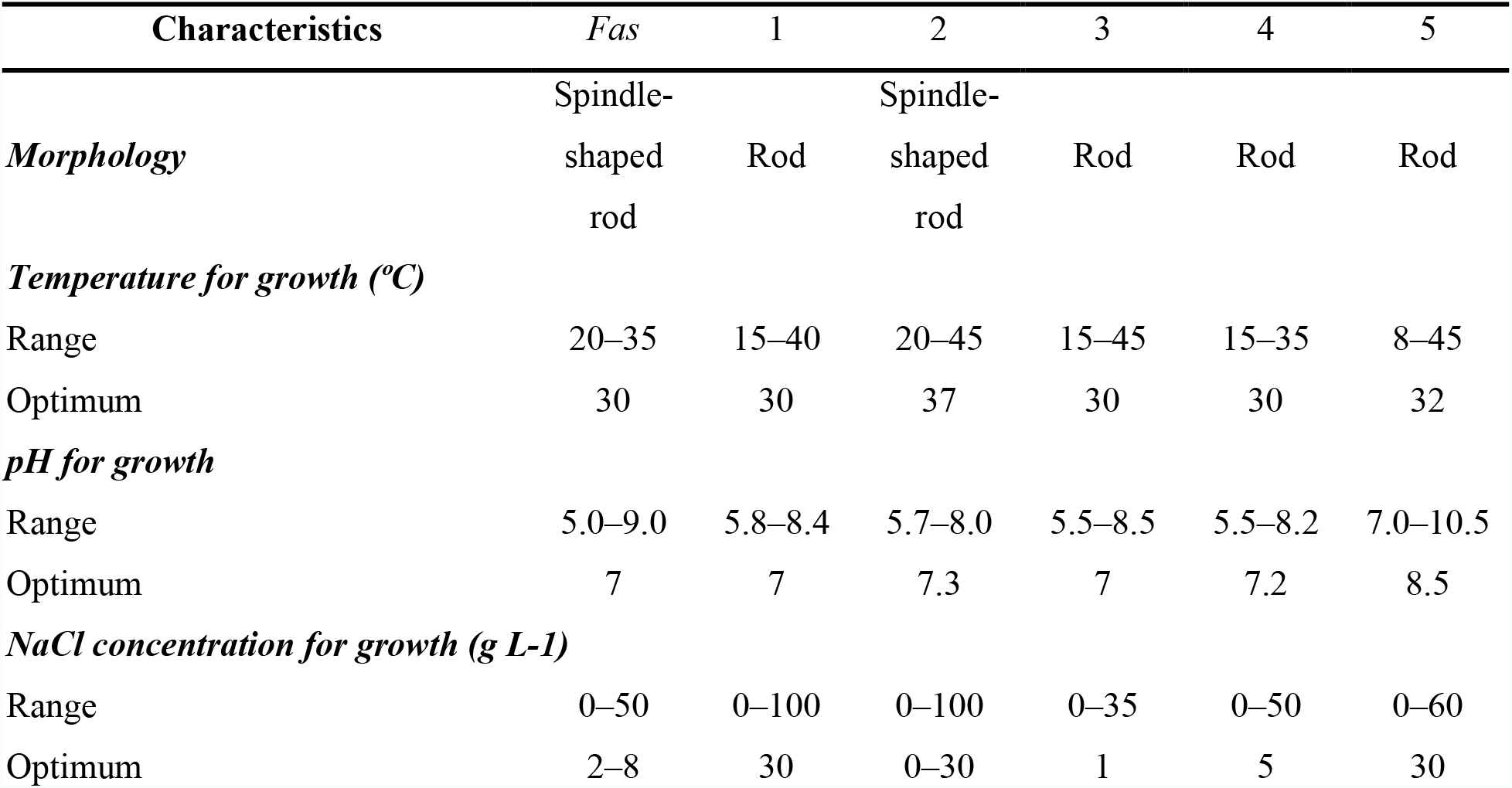
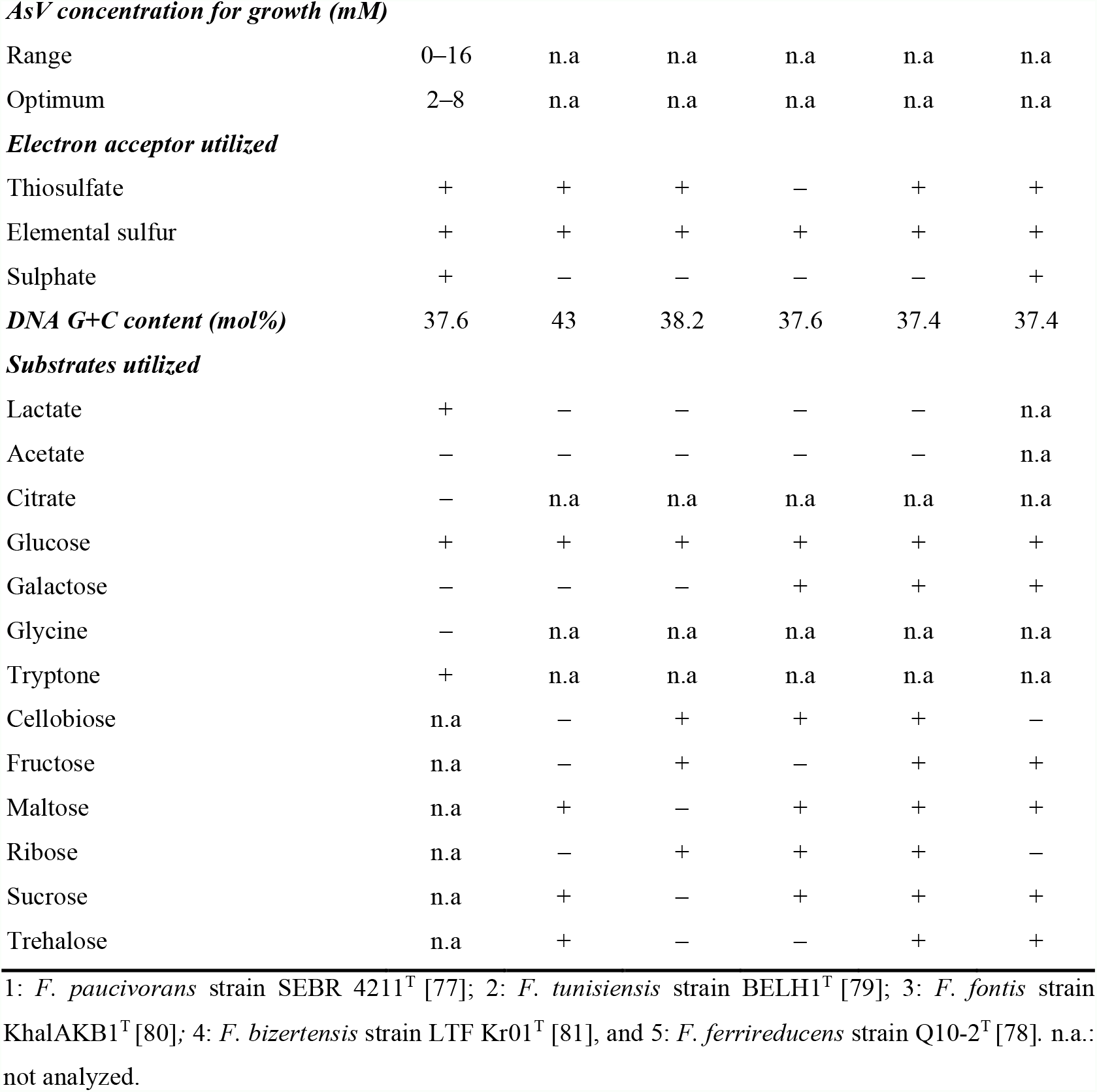
Characteristics that differentiate *Fusibacter* sp. strain 3D3 from other *Fusibacter* species.

**Figure 2.**
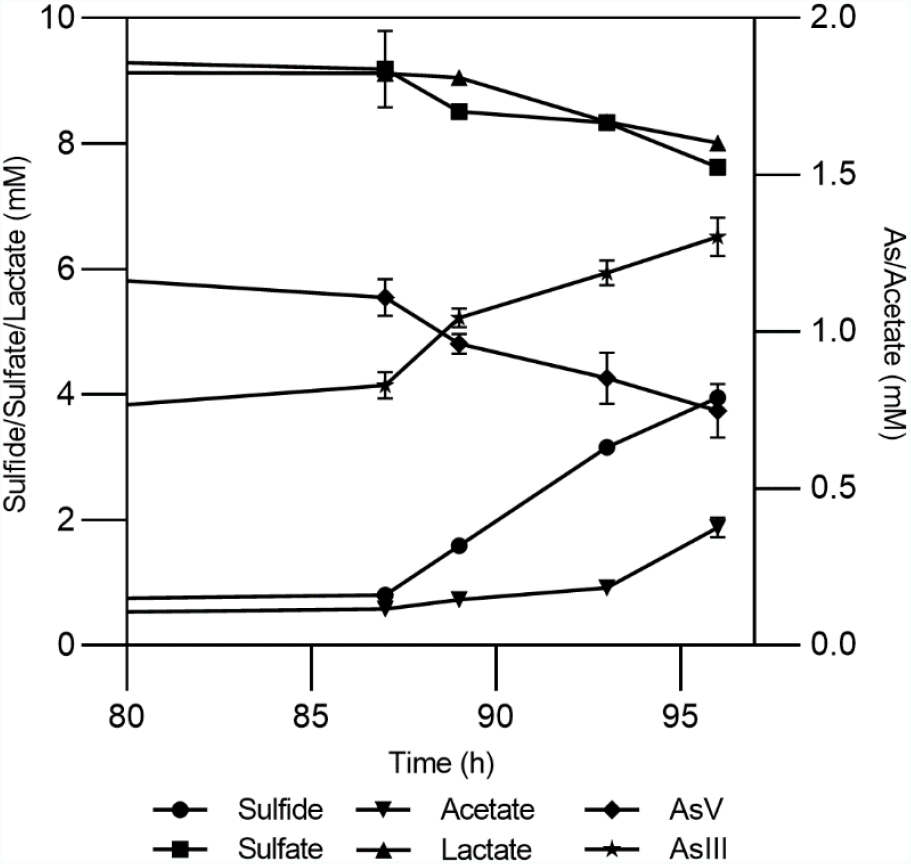
Growth of *Fas* in Newman-modified minimal medium with lactate (10 mM), sulfate (20 mM), arsenate (2 mM), yeast extract (0.1%), NaCl (10 g L-1), and cysteine (1 mM). Error bars represent the standard error of the mean of triplicate cultures. Sterile control experiments were also performed but the results were not shown for clarity.

*Fas* can be differentiated from *F. paucivorans, F. tunisiensis, F. fontis, F. bizertensis* and *F. ferrireducens* by its use of lactate as substrate, of sulfate as electron acceptor, the NaCl concentration for growth, its genomic DNA G+C content (Table 1) and its phylogeny [78]. Resistance to As was not reported for other isolated members of the genus.

Lactate was consumed (1.11 mM, with 0.26 mM of acetate formed) while arsenate (0.36 mM) and sulfate (1.56 mM) were reduced (Fig. 2). The amount of arsenite formed could not be determined quantitatively, as it tended to precipitate as yellow arsenic sulfide. On the other hand, sulfate reduction by *Fas* was demonstrated by sulfide and arsenic sulfide mineral production (Fig. 2). Interestingly, neither sulfate nor thiosulfate reduction was involved in energy conservation as it has been reported for other members of the *Fusibacter* genus [77, 79].

The ability of *Fas* to use lactate as electron donor when reducing arsenate suggests that arsenate respiration supports *its* growth. However, the AsV reduced/lactate oxidized molar ratio observed was 0.32±0.044 and the acetate produced/lactate consumed ratio was 0.23±0.05 (Fig. 2), when the theoretical values predicted for isolated arsenate reduction reactions when lactate is transformed to acetate by respiring microorganisms are 2 and 1, respectively [11]. In addition, the concomitant reduction of sulfate and the production of an arsenic sulfide precipitate does not allow an accurate quantification of arsenite and sulfide during *Fas* growth [13].

#### Assessment of specific sulfur species source for growth

The lack of differences in the sodium sulfate dose curve led us to study the role of sulfur sources in AsV reduction. *Fas* cultures with sodium sulfate, sodium thiosulfate and elemental sulfur were performed and combined with organic sulfur such as yeast extract and cysteine, both supplements required in Newman’s medium (Fig. S1). Culture without any source of inorganic sulfur was also carried out. All cultures were performed with 2 mM AsV.

The behavior of *Fas* cultures amended with sodium sulfate and sodium thiosulfate did not show significant differences, reaching the highest level of As reduction with Yeast/Cys complete medium. Furthermore, the intake of cysteine (empty square) as unique source of organic sulfur appeared to rise up to 50% of the total AsV reduction in all conditions at 96 hours, and it was especially evident when inorganic sulfur was absent. Cultures supplemented only with yeast extract (filled triangle) induced lower AsV reduction ratio than cysteine in all experiments. Growth (cell number) and sulfide production were also measured (Fig. S1).

#### Assessment of the role of proton/sodium gradient

The addition of 20 μM of sodium ion ionophore ETH2120 did not have a significant influence on the growth of the strain with lactate as the electron donor whether sulfate-arsenate (Fig. 3A squares) or only arsenate (Fig. 3B squares) were present as electron acceptors. On the other hand, 20 μM of the protonophore TCS completely inhibited the growth on lactate-sulfate-arsenate (Fig. 3A triangles) and lactate-arsenate (Fig. 3B triangles). Growth experiments showed that lactate-sulfate-arsenate and lactate-arsenate were insensitive to the Na^+^ ionophore ETH2120 but were highly sensitive to the protonophore TCS.

**Figure 3.**
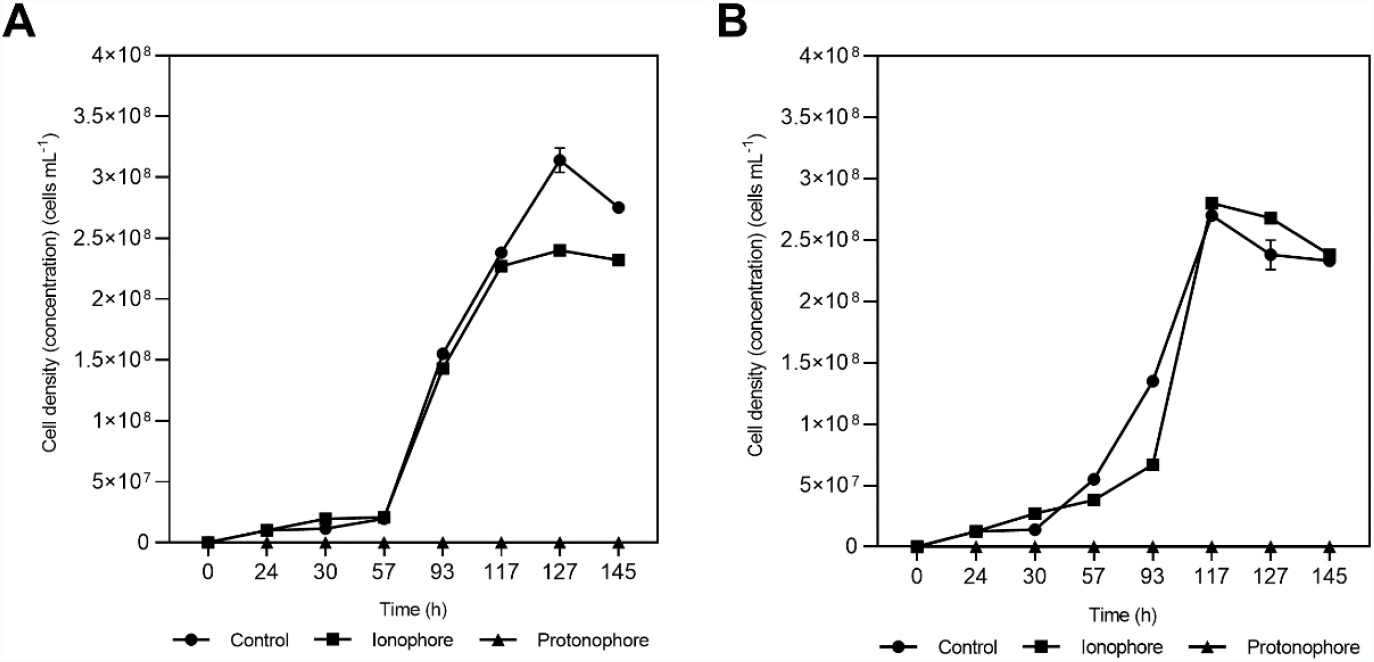
Effects of ETH2120 and TCS on *Fas* growth. Growth curves with lactate-sulfate-arsenate (A) and lactate-arsenate (B), with addition of 20 μM ETH2120 as ionophore (▪), 20 μM TCS as protonophore (▴) or no addition (•). Error bars show standard deviation of duplicates.

In the protonophore test, the resting cells were also decreasing and demonstrated to be highly sensitive to TCS suggesting that *Fas* needed the proton gradient for energy generation. Moreover, the inability of the strain to grow in the presence of TCS is consistent with the role of that gradient in the generation of a proton motive force.

#### Assessment of the Trx system in the response to As exposure

To gain insight into the thiol redox system involved in arsenate reduction and considering that the reductase ArsC of *Fas* was inferred by homology to be from the Trx/TrxR-dependent class, Trx (Fig. 4A) and TrxR (Fig. 4B) activity analysis in cellular extracts were performed. In addition, we compared the Trx activity with representative of Gram-positive (*B. subtilis*) and Gram-negative (*E. coli*) bacteria. The activity of Trx and TrxR increased after the exposure to As (Fig. 4).

**Figure 4.**
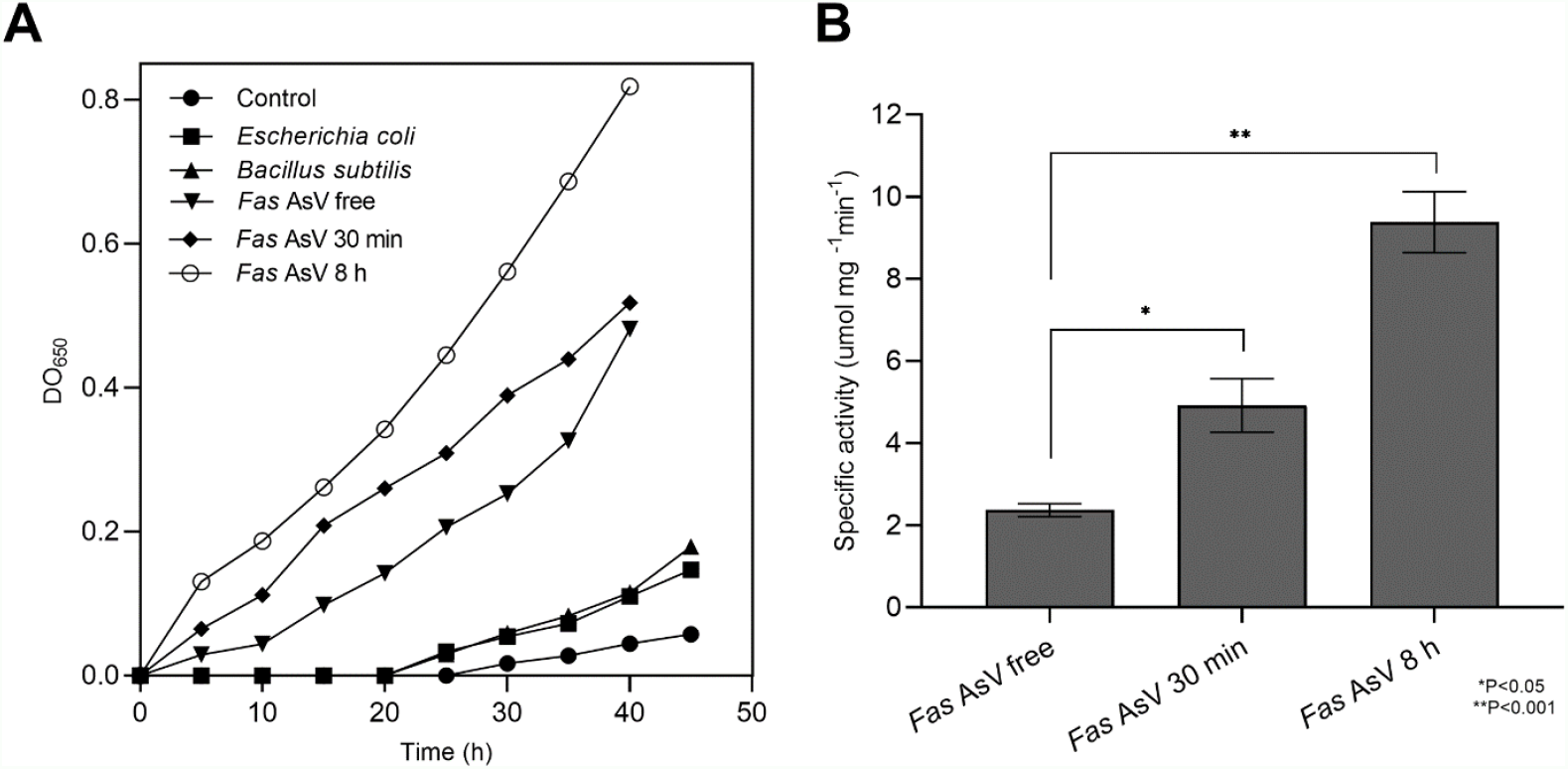
Measurement of Thioredoxin and Thioredoxin reductase activities. (A) Thioredoxin activity. The reduction of the insulin alfa-chain was monitored at 650 nm in 50 µg of whole cellular extract derived from *Fas* cells exposed to AsV during 30 min and 8 h and was compared to the activity in 50 µg of *Fas* cells grown without As and to other bacterial cultures. (B) Specific Thioredoxin reductase activity measured in a cell extract of *Fas* before and after AsV exposure.

### Genomic features

#### Genes involved in AsV reduction and energy metabolisms

As noted in Table 2, arsenic detoxification genes (*arsABCMR; acr3*) are clearly present in *Fas* genome [13]. *Fas* also contains genes coding for two putative cytoplasmic arsenate reductases with only 32% of identity in their aminoacidic sequences, and both clustered with genes coding for the thioredoxin-coupled family (Fig. S2). The revisited genomic context of *arsC*-2 (*arsD*-*arsR*-*pno*-*acr3*-*arsC*-2) [13] includes genes encoding for an arsenical resistant operon repressor (ArsD), a transcriptional regulator (ArsR), a 4Fe-4S ferredoxin (Pno, pyridine nucleotide-disulfide oxidoreductase NADH dehydrogenase), an arsenite efflux permease (Acr3), and the arsenate reductase (ArsC-2) [16] (Table 2). ATPase encoding gene that provide energy for AsIII efflux (*arsA*), included in the canonical *ars* operon of other Clostridiales, was also found in *Fas* even in another genomic context.

**Table 2.**
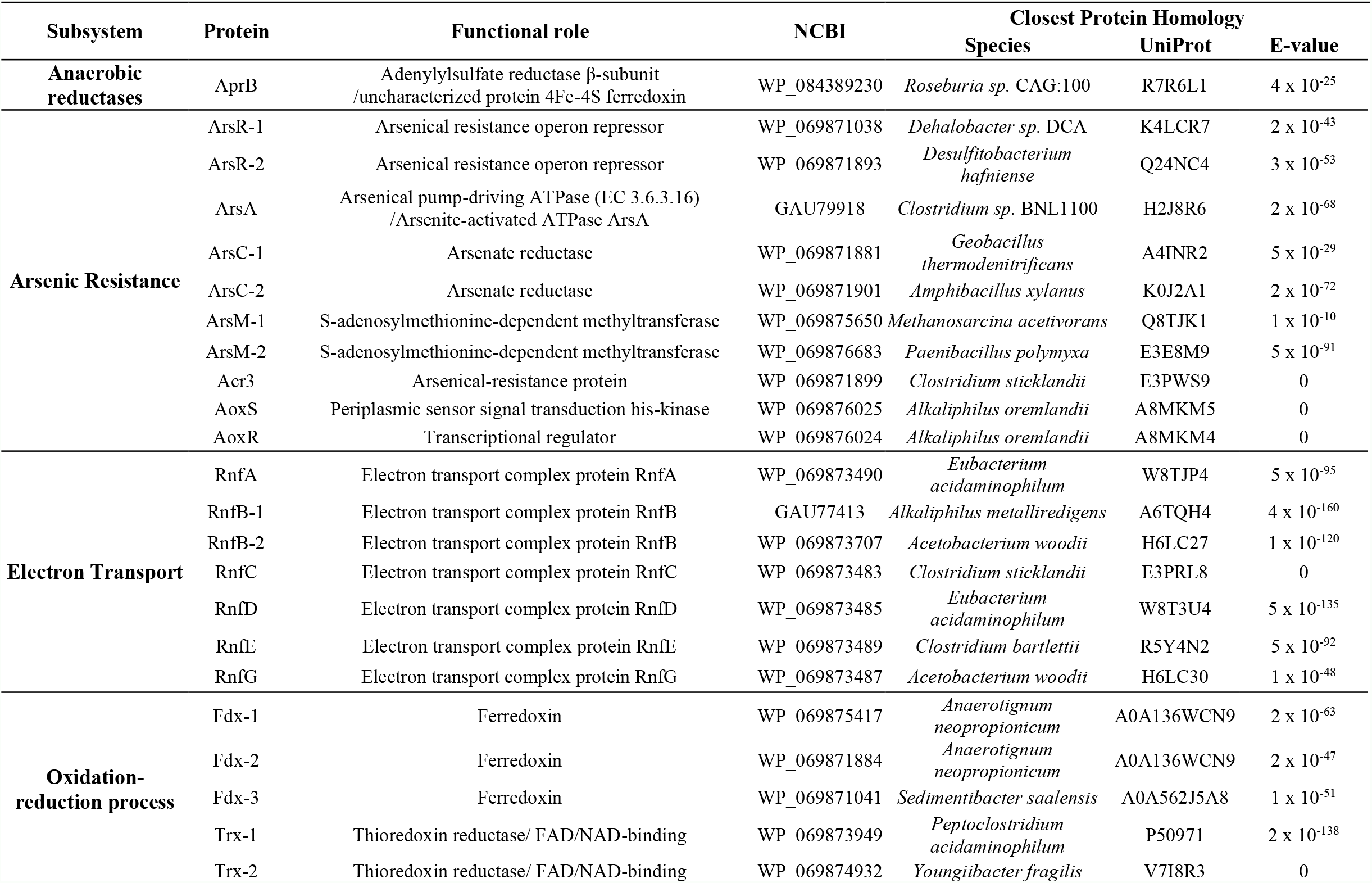
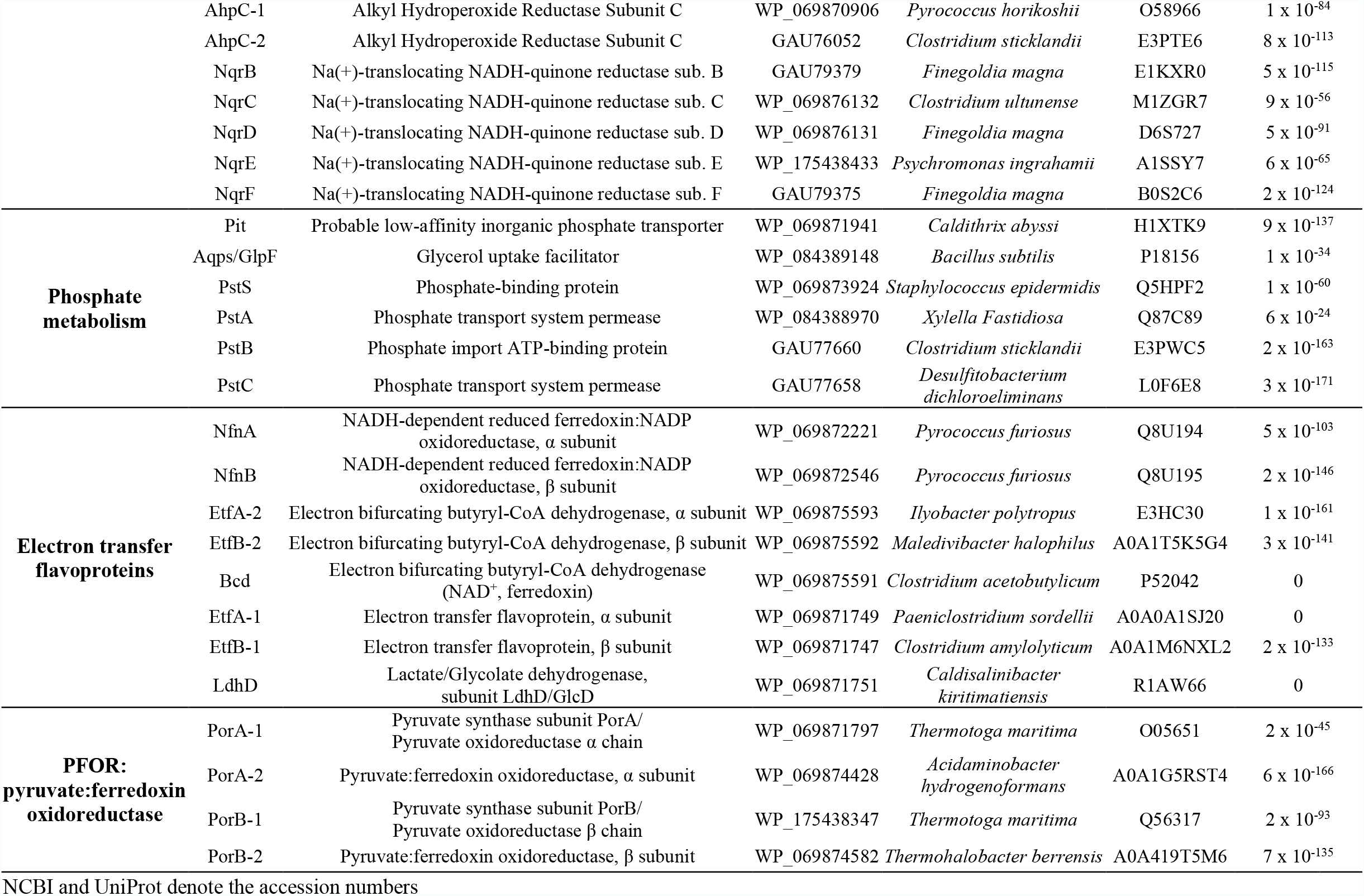
BLAST results of predicted proteins related to arsenic reduction and electron bifurcation in *Fas*.

By difference, the genomic context of *arsC*-1 revealed the presence of genes coding for ferredoxin and a redox-active disulfide protein (thioredoxin).

In other way, two thioredoxin reductase (TrxR) encoding genes were also identified in the *Fas* genome by the BLAST analysis (Table 2).

The dissimilatory arsenate reductase *arrAB* gene cluster, involved in anaerobic respiration using AsV as electron acceptor, was not found in *Fas* as it has been previously reported [13]. However, several genes predicted to be involved in the synthesis of the molybdenum cofactor included in the known catalytic site of ArrA [16] as well as in other cytoplasmic iron–sulfur proteins that catalyze ferredoxin-dependent redox reactions [34] were identified in the genome of *Fas*.

The NADH-dependent reduced ferredoxin:NADP oxidoreductase, α and β subunits (NfnAB) was evidenced by BLAST analysis against the *Pyrococcus furiosus* proteins [82] and it is conserved in the genomes of the *Fusibacter* genus. NfnAB is an electron bifurcating enzyme complex which couples the reduction of NADP+ with reduced ferredoxin (Fdred) and the reduction of NADP+ with NADH in a reversible reaction [50].

The constitutive *pit* (phosphate inorganic transport) and inducible *pst* (phosphate specific transport) operons involved in AsV uptake were also present in the *Fas* genome (Table 2).

The transmembrane ATP synthases (F0F1-ATPases) complex which is involved in ATP synthesis by obtaining the energy of a transmembrane gradient created by the difference in proton (H^+^) and in ATP hydrolysis in the reverse direction reactions are encoded in the *Fas* genome (Table S2). The order is conserved in the genomes of the *Fusibacter* genus (subunits *I, A, C, C, B*, Delta, Gamma, Beta, Epsilon).

In addition, the occurrence of the genes *rnfC, D, G, E, A, B* reported as encoding for the membrane-associated ferredoxin-dependent *Rhodobacter* nitrogen fixing (Rnf) complex responsible for transmembrane Na^+^/H^+^ transport [40] and for Na^+^/H^+^ gradient harvesting [36] was revealed by BLAST analysis in *Fas* genome, and in the whole genus (Table 2).

A search for genes involved in the fermentation process [83] in *Fusibacter* genomes revealed the occurrence of genes coding for an aldehyde dehydrogenase and butanoate metabolism in most of them, while genes involved in lactate/pyruvate metabolism were not present neither in *F. tunisiensis* nor in *F. paucivorans*. Genes codifying for pyruvate decarboxylase, alcohol dehydrogenase (cytochrome c), and proteins involved in the citrate cycle were absent (Table 3).

**Table 3.**
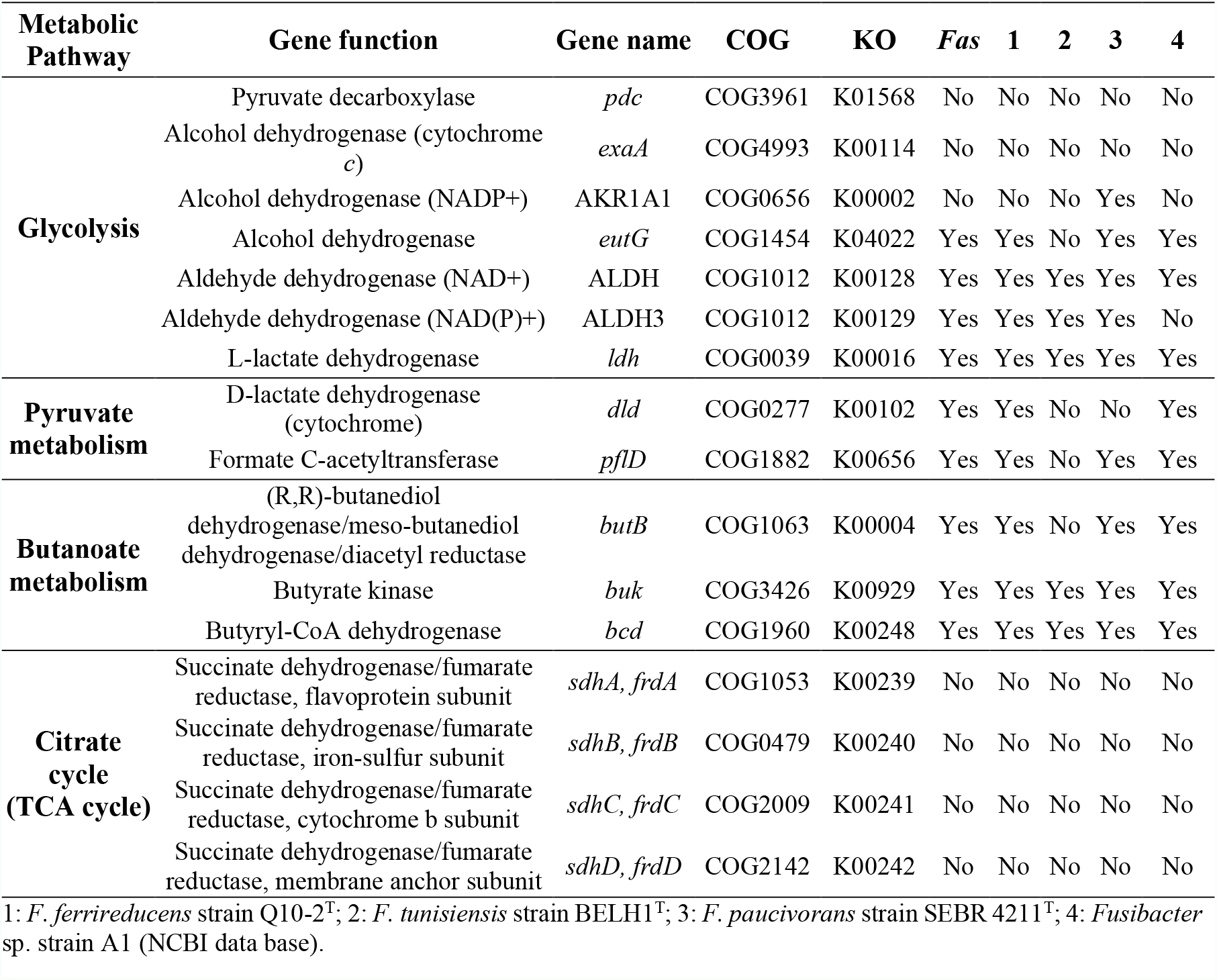
Presence of genes related to fermentative metabolism in *Fusibacter* genomes.

Searching for *etfB* genes in the *Fusibacter* genomes allowed us the finding of genomic contexts that would code for proteins involved in electron bifurcation. To verify the presence of key features (motifs 1 and 2) of the electron bifurcating EtfBs, an alignment of putative *Fusibacter* EtfBs with previously characterized proteins was performed (Fig. 5). We found that all *Fusibacter* genomes code for group 2A EtfBs, *Fas* and *F. ferrireducens* also code for group 2B, while only *F. paucivorans* code for group 2C elements. In group 2A, all the putative proteins from *Fusibacter* process the key conserved residues. In group 2B, the Thr-94 is not conserved and is substituted by a valine in *Fas* and *F. ferrireducens* EtfB-1.

**Figure 5.**
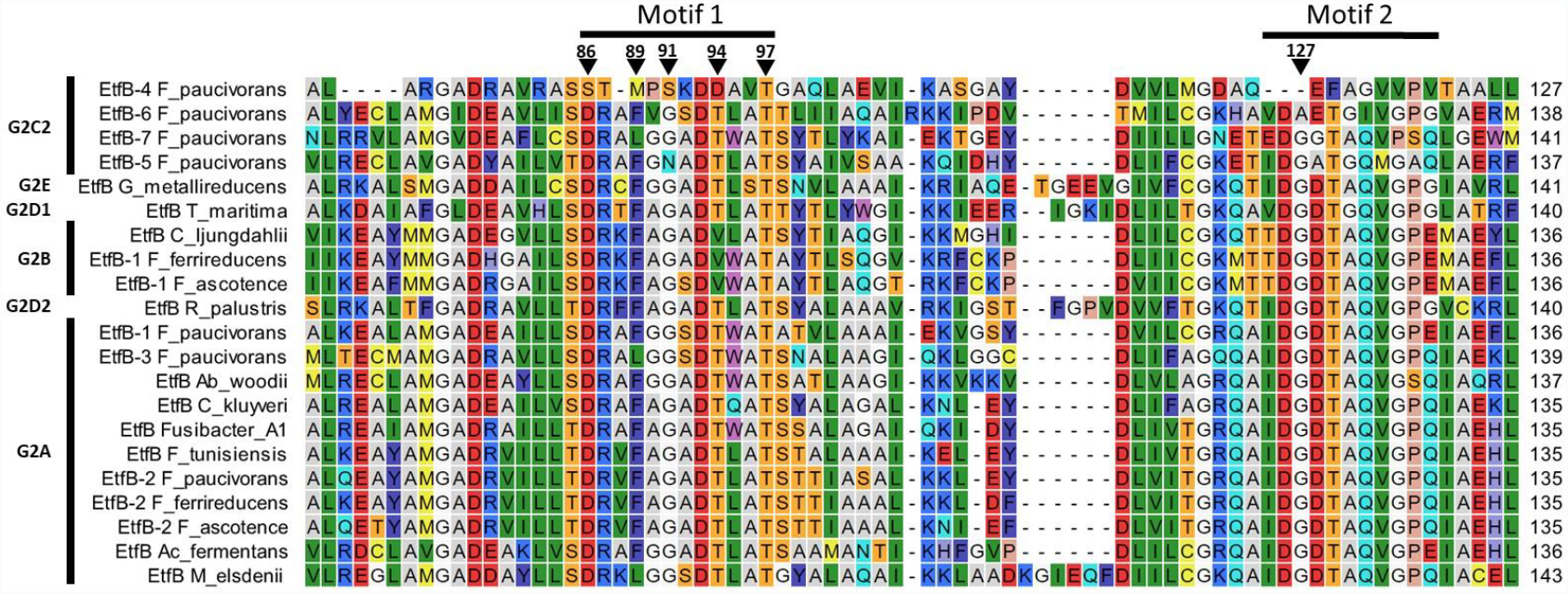
Protein sequence alignment to compare characterized electron-transferring flavoproteins with those from *Fusibacter*. The horizontal bars indicate motifs 1 and 2, corresponding to the NADH-and FAD-binding sites in bifurcating Etfs, respectively. The inverted triangles indicate residues proposed to coordinate NADH and FAD and the numeration corresponds to EtfB *R_palustris*. The Etf groups are indicated on the left part. Representatives of EtfB groups are from *Geobacter metallireducens* GS-15 (*G_metallireducens*; YP_383650), *Thermotoga maritima* MSB8 (*T_maritima*; NP_229330), *Clostridium ljungdahlii* PETCPETC (*C_ljungdahlii*; YP_003780321), *Rhodopseudomonas palustris* BisA53 (*R_palustris*; YP_783418), *Acetobacterium woodii* WB1 (*Ab_woodii*; AFA48355), *Clostridium kluyveri* DSM 555 (*C_kluyveri*; YP_001393858), *Acidaminococcus fermentans* VR4 (*Ac_fermentans*; YP_003398269), *Megasphaera elsdenii* T81 (*M_elsdenii*; WP_022498188). *Fusibacter* proteins are listed in Table S3.

To better understand their possible role in *Fas*, we compared the genomic contexts of the *etf* encoding genes in *Fusibacter* (Fig. 6). All the analyzed genomes contain the gene encoding for the electron transfer flavoprotein subunit beta followed by the alpha subunit encoding gene. We found at least one copy of the genes encoding for EtfA, EtfB, and a putative butyryl-CoA dehydrogenase (Bcd) in *Fusibacter* genomes. Interestingly, *bcd* is always located upstream of *etf* genes cluster.

**Figure 6.**
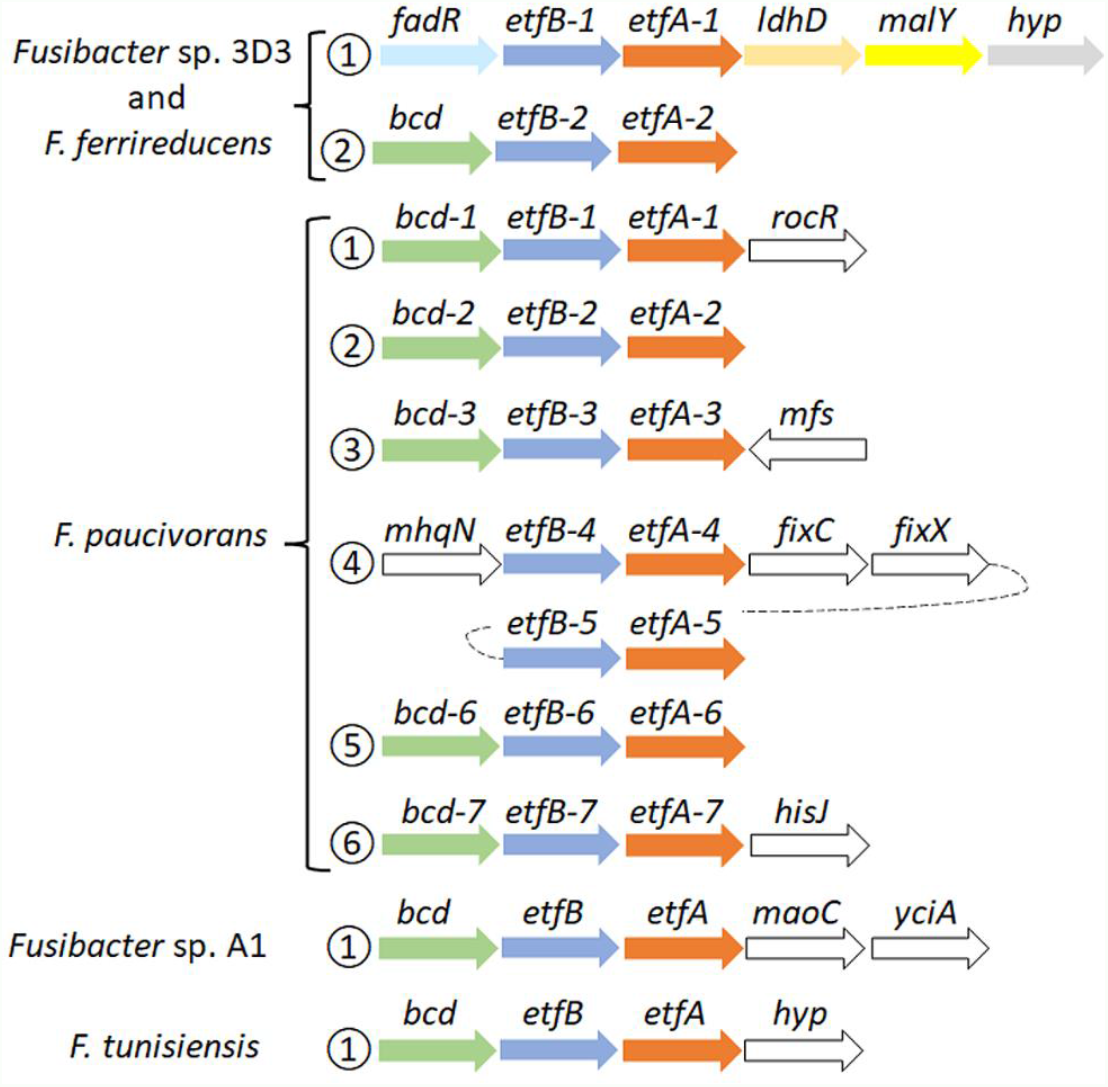
Genomic context of *etf* related genes in *Fusibacter*. Numbers in circles indicate the occurrence of the *etf* copies. The arrows represent the orientation of the gene (size is not at scale), and the same colors indicate homology, except white. The gene products are described in the text and the gene product accession codes are listed in Table S3.

*Fas* and *F. ferrireducens* have two identical genetic arrangement. In context 1, *fadR* (which codifies for a transcriptional regulator) is located upstream *etfB-1* and *etfA-1* genes and downstream of both are *ldh* that codifies for a Lactate/Glycolate dehydrogenase (COG0277), *malY* that codifies a putative pyridoxal 5’-phosphate (PLP)-dependent C-S lyase (COG1168) and a gene that codifies for a hypothetical protein conserved in both genomes, while in context 2, only a *bcd* gene was identified upstream *etfBA-2* (Fig. 6).

Surprisingly, *F. paucivorans* contains seven copies of the *etfB-etfA* pair in six different contexts. Contexts 1, 2, 3, and 6 present the upstream arrangement with the *bcd* gene (Fig. 6). In context 4, *mhqN*, which codifies for a nitroreductase family protein (cd02137), is found upstream *etfBA-4* while *fixC* and *fixX*, which code for a flavoprotein dehydrogenase (COG0644) and a ferredoxin-like protein (COG2440) respectively.The fifth copy *etfBA-5* was identified downstream *fixX* (Fig. 6). In addition, *F. paucivorans* has four orphan *etfB* genes, possibly belonging to the 2C2 group due to its phylogeny and because it does not have any *etfA* or *etfB* genes fused in a single open-reading frame, as it has been described in the group 2C1 [75]. Independently, we also identified in *F. paucivorans* genes coding for a nitrogenase reductase and maturation protein (*nifH*), the regulatory proteins P-II (*glnA* and *glnB*) and α and β subunits of the nitrogenase (*nifD* and *nifK*). This opens the possibility that in this *Fusibacter* species some *etf* genes participate in nitrogen fixation. Indeed, no other *Fusibacter* possesses nitrogen fixation genes (results not shown).

*Fusibacter* sp. A1 and *F. tunisiensis* had only one specific genomic context with *etf*-related genes. In *Fusibacter* sp. A1, downstream *etfA* we found *maoC*, that codifies for an acyl dehydratase (COG2030), followed by *yciA*, coding for an acyl-CoA hydrolase (COG1607), both related to lipid transport and metabolism (Fig. 6).

Finally, the subsystem approach to genome annotation performed by RAST/SEED [67] confirmed the relatedness of *Fas* to other members in the Clostridiales order (Table 2).

#### Detection of dissimilatory arsenate reductase *arrAB* genes

The dissimilatory arsenate reductase *arrAB* gene cluster, involved in anaerobic respiration using AsV as electron acceptor, was not found in *Fas* (Fig. S3). This suggests that a different mechanism independent of ArrAB is conferring the ability to obtain energy from AsV reduction.

#### Heterologous expression of *arsC* genes

Two *Fas* arsenate reductases encoding genes were identified in the genome sequence and specific primers were designed for their amplification from *Fas* genomic DNA by PCR (Fig. S4). The amplified genes (*arsC*-1_*Fas*_ and *arsC*-2_*Fas*_) were first cloned in the pGEM-T cloning vector, then released through enzymatic DNA digestion (Fig. S5) and ligated into the pTrcHis2A expression vector. The presence of the insert in the expression vector was checked by colony PCR (Fig. S6) or releasing the insert through plasmidic DNA digestion (Fig. S7). The activity of the gene product coded by the insert was tested by growing the recombinant *E. coli* WC3110 in the presence of AsV (Fig. 7). Complementation of the Δ*arsC E. coli* WC3110 strain with the insert of both putative *arsC*_*Fas*_ genes evidenced changes in AsV. A higher resistance to AsV was conferred by ArsC-2_*Fas*_ compared to ArsC-1_*Fas*_ (Fig 7). Growth of *E. coli* WC3110 strain without insert was not observed. These biological data are the first metabolic evidence needed to confirm the existence of the proposed metabolism in *Fas*.

**Figure 7.**
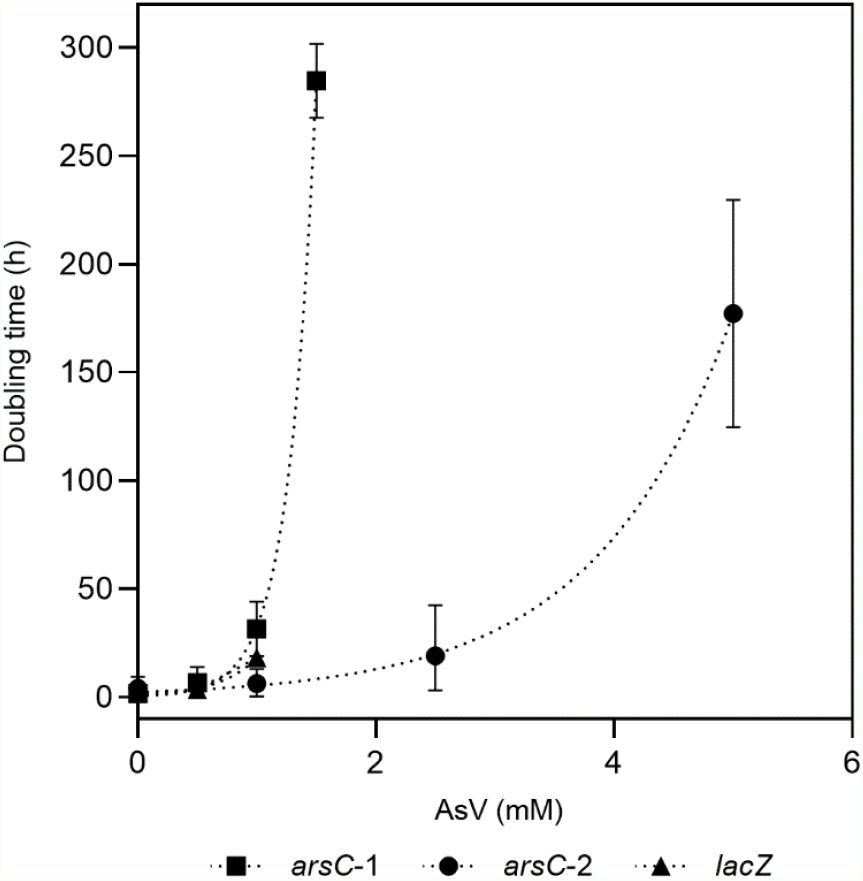
Evaluation of resistance to AsV conferred by *arsC*-1 and *arsC*-2 from Fas. Growth of *ΔarsC E. coli* WC3110 strain complemented by *arsC*-1_*Fas*_ SHT, *arsC*-2_*Fas*_ SHT or *lacZ* genes in presence of AsV.

## Discussion

Phylogenetic analysis performed with the 16S rRNA genes had formerly grouped *Fas* inside the Gram-positive *Fusibacter* genus [13]. The *in silico* average nucleotide identity (ANI) with its closest relative type strain is 80.1% [78] which allows the confirmation that *Fas* affiliates with the *Fusibacter* genus (> 70%) [84], and support the proposal of *Fusibacter ascotence* as a new species of the genus (<95-96%).

Taking together the observed growth features of *Fas* compared with other species of the *Fusibacter* genus (Table 1), and the insights into their genome sequences (Tables 2, 3 and S1) we are able to propose a rationale to justify the singularity of the *Fas* energetic metabolism.

- It grows strictly in anaerobiosis by reducing arsenate and using lactate as electron donor and its growth is improved by increasing AsV concentration (Fig. 1), being 2 mM the optimum level. To date, arsenic metabolism was not reported for the other *Fusibacter* species.
- Despite the arsenic reducing activity, the dissimilatory arsenate reductase *arrA* gene was not detected neither by the *Fas* genome sequence analysis [13] nor by PCR assays (Fig. S3). Moreover, neither *arrC* (coding for the membranous subunit suggested to play the role of menaquinone oxidation) reported to be present in some AsV reducing bacteria [8], nor *omc* (encoding an outer-surface, octaheme c-type cytochrome), and *cymA* (encoding a membrane-attached MKH2 oxidizing protein) genes reported in arsenic respiring *Shewanella* sp. strains [9, 85] were evidenced in the *Fas* genome (Table 2). The heterologous expression on *ΔarsC E. coli* WC3110 strain has allowed us to confirm that ArsC-1 and ArsC-2 of *Fas* are functional and confer As resistance (Fig. 7). Both *arsC*_*Fas*_ genes belong to the Enterobacterial clade one [20], and therefore encode a TrxR-dependent class of ArsC. In addition, the enzymatic analysis revealed a high Trx and TrxR activity in cells cultured with As, supporting the inference about the Trx dependence of the ArsC in *Fas*.
- All the previously reported strains inside the *Fusibacter* genus are fermentative bacteria [77–81]. Interestingly, *Fas* can use lactate and glucose as substrates, while. *F. tunisiensis, F. paucivorans, F. bizertensis*, and *F. ferrireducens* can not utilize lactate [77–79, 81]. The genetic evidence agrees with the observed physiology on the culture conditions tested (Table 3).
- Furthermore, all the reported *Fusibacter* species have the ability to reduce sulfured nutriments [77–81]. Sulfate reduction by *Fas* was demonstrated by sulfide and arsenic sulfide mineral production, and thiosulfate reduction was also positively checked. In addition, thiosulfate and sulfate were more efficient than S° to stimulate cell growth (Fig. S1) perhaps because of the low solubility of S°. Other *Fusibacter* species reduce thiosulfate and sulfur (but not sulfate or sulfite), and only *F. ferrireducens* shares with *Fas* the ability to reduce sulfate. Neither sulfate nor thiosulfate were involved in energy conservation in *Fas* as it has been reported for the other members of the *Fusibacter* genus [79, 80]. That feature could also be related to other cellular mechanisms present in microorganisms to cope with stress, such as arsenic stress, i.e. sulfur assimilation [86]. *F. paucivorans* growth experiments with sulfured nutriments revealed that the addition of thiosulfate relieved the inhibition produced by the H_2_ released by the glucose fermenting metabolism. In addition, a differential pattern of glucose fermentation products was observed in cultures with thiosulfate, represented by a decrease in butyrate levels together with an increase in acetate production [77]. As well as for other fermenting bacteria, those results confirm the Huber hypothesis that sulfur reduction plays a role of an electron sink reaction to prevent H_2_ accumulation from fermentation metabolism [87]. Therefore, the lactate/butyrate fermentation metabolism in *Fusibacter* should be regulated by the cellular redox state resembling the reported for other *Firmicutes* [88]. Interestingly a redox-sensing transcriptional repressor gene encoding a protein whose DNA binding activity is modulated by the NADH/NAD^+^ ratio [88] is located downstream to the Acetyl-CoA acetyltransferase encoding gene in *Fas* and other *Fusibacter* genomes (data not shown). The Acetyl-CoA acetyltransferase is in charge of the first step during the Acetyl-CoA fermentation to Butyrate pathway after the split in the three alternative fermentation pathways.
- The results obtained from the growth experiments with and without addition of the protonophore TCS and the ionophore ETH2120 [41] revealed that *Fas* does require the formation of a proton gradient to get energy for growing on AsV (Fig. 3). In addition, the occurrence of genes encoding for the Rnf complex in *Fas* genome (Table 2) allows us to infer the capacity of *Fas* for energy conservation/utilization via proton translocating ferredoxin oxidation/reduction. Finally, the F0F1ATP synthase would couple ATP synthesis to the electrochemical gradient based on differences in the proton concentration generated.
- In agreement with the genomic characterization of the ArsC_*Fas*_ inside the Trx-dependent class, the enzymatic analysis has shown an increased level of Trx and TrxR activities after AsV addition (Fig. 4). Besides, it is known that thioredoxin is also involved in sulfur assimilation evidenced in the early response to arsenic and in maintaining the cellular redox state [86].
- *Fas* has all the known genomic resources for the pathway of lactate fermentation to acetate and butyrate in *Firmicutes* [88]. Interestingly, the genomes of *Fas* and *F. ferrireducens* contain two genomic contexts that may be involved in the electron bifurcation process of the electron-transferring flavoproteins (EtfAB) type [50]. According to the model proposed [88] for lactate and acetate transformation to butyrate, there must be a lactate dehydrogenase/EtfAB complex and a butyryl CoA dehydrogenase/EtfAB complex (Fig. 8A). We propose that contexts 1 and 2 encode the elements for the transformation of lactate and butyryl CoA, respectively (Fig. 6). Acetate production was observed in *Fas* and butyrate plus acetate production was confirmed in *F. paucivorans* [77]. *Clostridium butyricum* and *Acetobacterium woodii* were shown to transform lactate during fermentation by an enzyme complex of Ldh, EtfAB [57, 88]. Analysis of the genomes of *C. butyricum*, and *A. woodii* among others, showed conservation of the genetic context of at least the genes coding for these proteins [52, 57, 88]. Proteins related to butyryl CoA transformation involve butyryl-CoA dehydrogenase/electron transfer flavoproteins EtfA and EtfB [89]. This complex is also encoded in a conserved array [52, 88]), like the genomic context 2 found in *Fas* (Fig. 6).
- Furthermore, the occurrence of a NADH dependent reduced ferredoxin NADP+ oxidoreductase complex, the second type of FBEB complexes [50] in *Fas* genome (Table 2), permits to hypothesize that energy for growth should be provided by the energetic link of cellular ferredoxin and NAD^+^ pools through the Rnf function to generate chemiosmotic potential when ferredoxin is higher than NADH level or, in reverse, for ferredoxin generation when NADH is higher [90], for a more efficient metabolism in anoxic environments. In addition, Nfn could play the reported role of balancing the redox state of the pyridine nucleotide NAD(H) and NADP(H) pools and, in that way, favor the catabolic or anabolic reactions [52].
- The occurrence of multiple and different bifurcating (Bf) enzymes observed in *Fas* has been already detected in several *Firmicutes* genomes, and Bf-Ldh, Bf-Bcd and Nfn, that share NAD(H) and ferredoxin as common substrates, usually participate in those combinations [82].

**Figure 8.**
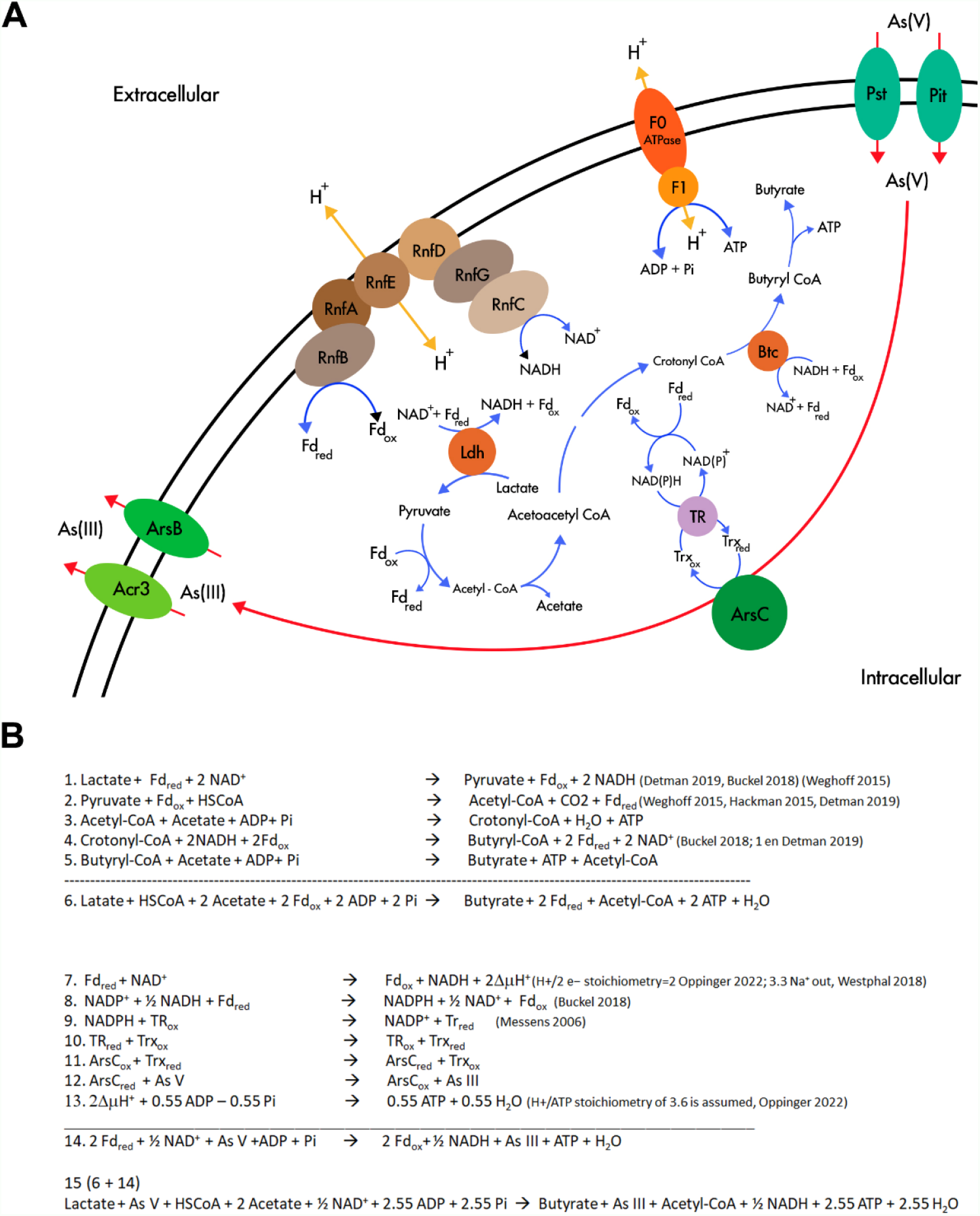
Flavin based electron bifurcation and its link with AsV reduction. (A) Working model of the involvement of FBEB and the AsV reduction in *Fas*. (B) Stoichiometry of the metabolism proposed in (A).

The revealed specialization for lactate fermentative metabolism present in *Fas* and already reported in *Firmicutes* [88] with the participation of FBEB and Rnf supports the availability of NADH and Fd_red_ and ATP generation (Figure 8 and Equations 1-5 in Fig. 8B). NADH and Fd_red_ should be the soluble electron carriers required for producing NADPH and starting the cascade of thiol reductases (NADPH→TR→Trx→ArsC) [29] involved in AsV reduction by ArsC Trx type of *Fas* (Fig. 8A and Fig. 8B, equations 7-15). In that way, besides to generate ΔμH+ coupled to ATP synthesis by ATP synthase, the FBEB would conduct the reduction of AsV by providing the low potential ferredoxin. The AsIII efflux pumps present in the Ars operon allow AsIII elimination and As_4_S_4_ precipitation outside the cells.

The analysis of the reported stoichiometry [29, 37, 50, 57, 90] hints us that it is plausible that AsV could play a role similar to CO_2_ in acetogenic bacteria [39], of terminal acceptor of the electrons derived from reduced ferredoxin, the low potential electron carrier generated by electron bifurcation (Fig. 8B).

This rationale allows us to formulate a hypothetic metabolism (Fig. 8A) similar to the evidenced in other anaerobic microorganisms [51]: Arsenate reduction provides additional energy to arsenic reducing fermenters independent of ArrAB for growing through a new mechanism that involves soluble ferredoxin electron carrier, FBEB complexes, the cytoplasmic ArsC, and the membrane-associated ion-translocating complex Rnf. As previously reported, this system could be regulated by the redox state [88]. The energetic link of cellular NADH and ferredoxin should be the way in which the electrons reach AsV in the cytoplasm, converting it in an electron sink/electron acceptor, similar to the role assigned to ferric iron in *F. ferrireducens* [78].

Finally, metagenomic analysis evidenced that the Trx-ArsC is much more diverse in the high altitude modern stromatolites in the Argentinian Puna (Altiplano), than at the base of the Socompa Volcano [91] characterized by high arsenic contents. Furthermore, the ecological relevance of the proposed metabolism was suggested by the dominance of genes predicted for encoding the Trx-ArsC versus Grx-ArsC cytoplasmic arsenate reductase in arsenic rich environments on a regional survey at the High Andes [31] where *Fas* was isolated from.

## Acknowledgements

The authors are appreciative of the technical support of Olga Encalada, Dr. Antonio Serrano, Dr. Lorena Escudero, Dr.(C) Cinthya Tebes-Cayo, Dr. Nia Oetiker, and of María Celia Chong D. for her support in the model designing.

