## Supplemental Figures and Tables for "Looking for the mechanism of arsenate respiration in an arsenate-dependent growing culture of *Fusibacter* sp. strain 3D3, independent of ArrAB"

**Table S1. Primers used to clone *arsC*-1 and *arsC*-2 gene from *Fas*.**

| **Primers** | **Sequences 5'-3'** | **Annealing (°C)** |
| --- | --- | --- |
| arsC-1_*Hin*dIII_Rv | aagcttTCAATCTAAATTGTATTTGCTCTT | 56.1 |
| arsC-1_SHT_*Xho*I_Fw | ctcgagTatGAAGCCCGTTAAAATTTTA |  |
| arsC-1_HT_*Xho*I_Fw | ctcgagATGAAGCCCGTTAAAATTTTA |  |
| arsC-2_*Hin*dIII_Rv | aagcttTTATAAATCAAGCACAATTTG | 60.0 |
| arsC-2_SHT_*Xho*I_Fw | ctcgagtATGAGTAGAAAACCAAAAG |  |
| arsC-2_HT_*Xho*I_Fw | ctcgagATGAGTAGAAAACCAAAAG |  |
| Lowercase letters indicate changes made from the original sequence | | |

**Table S2. Presence of genes related to ATP synthesis.**

| **NCBI** | **Function** | **Closest protein homology** | | | |
| --- | --- | --- | --- | --- | --- |
|  |  | **Identity** | **E-value** | **UniProt** | **Organism** |
| WP_069870490 | F0F1 ATP synthase subunit epsilon (atpC) | 51.4% | 5.9x10^-39^ | A8MJV8 | *Alkaliphilus oremlandii* OhILAs |
| WP_069870492 | F0F1 ATP synthase subunit beta (atpD) | 76.9% | 0.0 | A8MJV9 | *Alkaliphilus oremlandii* OhILAs |
| WP_069870494 | ATP synthase F1 subunit gamma (atpG) | 53.6% | 2.3x10^-99^ | [A6TK64](https://www.uniprot.org/uniprot/A6TK64) | *Alkaliphilus metalliredigens* QYMF |
| WP_069870496 | F0F1 ATP synthase subunit alpha (atpA) | 75.5% | 0.0 | A8MJW1 | *Alkaliphilus oremlandii* OhILAs |
| WP_069870498 | F0F1 ATP synthase subunit delta (atpH) | 37.8% | 3.8x10^-35^ | A8MJW2 | *Alkaliphilus oremlandii* OhILAs |
| WP_069870500 | F0F1 ATP synthase subunit B (atpF) | 48.4% | 1.1x10^-46^ | Q0ZS23 | *Clostridium paradoxum* |
| WP_069870502 | ATP synthase F0 subunit C  (atpE) | 86.4% | 2.7x10^-45^ | A8MJW4 | *Alkaliphilus oremlandii* OhILAs |
| WP_069870504 | ATP synthase F0 subunit C  (atpE) | 71.4% | 2.6x10^-34^ | A8MJW4 | *Alkaliphilus oremlandii* OhILAs |
| WP_242877057 | F0F1 ATP synthase subunit A (atpB) | only unreviewed entries available | | | |
| WP_069870509 | ATP synthase subunit I  (atpI) | only unreviewed entries available | | | |
| Accession numbers to databases NCBI (*Fusibacter* sp. strain 3D3 genes) and UniProtKB (closest reviewed entry) are under NCBI and UniProt columns, respectively. | | | | | |

**Table S3. Products of the genes in the genetic contexts of Etfs in *Fusibacter*.** The *etf* gene contexts, EtfB group and NCBI accession number are indicated.

| **Microorganisms** | **Context** | **Group** |  |  |  |  |  |  |  |
| --- | --- | --- | --- | --- | --- | --- | --- | --- | --- |
| ***Fas*** | **1** | **G2B** | **Proteins** | **FadR** | **EtfB-1** | **EtfA-1** | **Ldh** | **MalY** | **Hyp1** |
|  |  |  | **Accession number** | WP_069871746 | WP_069871747 | WP_069871749 | WP_069871751 | WP_069871752 | WP_069871754 |
|  | **2** | **G2A** | **Proteins** | **Bcd** | **EtfB-2** | **EtfA-2** |  |  |  |
|  |  |  | **Accession number** | WP_069875591 | WP_069875592 | WP_069875593 |  |  |  |
| ***F. ferrireducens*** | **1** | **G2B** | **Proteins** | **FadR** | **EtfB-1** | **EtfA-1** | **Ldh** | **MalY** | **Hyp1** |
|  |  |  | **Accession number** | WP_194701605 | WP_194701606 | WP_194701607 | WP_194701608 | WP_194701609 | WP_194701610 |
|  | **2** | **G2A** | **Proteins** | **Bcd** | **EtfB-2** | **EtfA-2** |  |  |  |
|  |  |  | **Accession number** | WP_194702816 | WP_194702815 | WP_194702814 |  |  |  |
| ***F. paucivorans*** | **1** | **G2A** | **Proteins** | **Bcd-1** | **EtfB-1** | **EtfA-1** | **RocR** |  |  |
|  |  |  | **Accession number** | WP_213235038 | WP_213235037 | WP_213235036 | WP_213235035 |  |  |
|  | **2** | **G2A** | **Proteins** | **Acd-2** | **EtfB-2** | **EtfA-2** |  |  |  |
|  |  |  | **Accession number** | WP_213235641 | WP_213235640 | WP_213235639 |  |  |  |
|  | **3** | **G2A** | **Proteins** | **Acd-3** | **EtfB-3** | **EtfA-3** | **Mfs** |  |  |
|  |  |  | **Accession number** | WP_213236303 | WP_213236302 | WP_213236301 | WP_213236300 |  |  |
|  | **4** | **G2C2** | **Proteins** | **MhqN** | **EtfB-4** | **EtfA-4** | **FixC** | **FixX** |  |
|  |  |  | **Accession number** | WP_213238184 | WP_213238183 | WP_213238182 | WP_213238181 | WP_213238180 |  |
|  |  | **G2C2** | **Proteins** |  | **EtfB-5** | **EtfA-5** |  |  |  |
|  |  |  | **Accession number** |  | WP_213238179 | WP_213238178 |  |  |  |
|  | **5** | **G2C2** | **Proteins** | **Acd** | **EtfB-6** | **EtfA-6** |  |  |  |
|  |  |  | **Accession number** | WP_213238432 | WP_213238433 | WP_213238434 |  |  |  |
|  | **6** | **G2C2** | **Proteins** | **Acd** | **EtfB-7** | **EtfA-7** | **HisJ** |  |  |
|  |  |  | **Accession number** | WP_213238473 | WP_213238472 | WP_213238471 | WP_213238470 |  |  |
| ***Fusibacter* sp. A1** | **1** | **G2A** | **Proteins** | **Acd** | **EtfB** | **EtfA** | **MaoC** | **YciA** |  |
|  |  |  | **Accession number** | WP_129487661 | WP_129487660 | WP_129487659 | WP_129487658 | WP_129487657 |  |
| ***F. tunisiensis*** | **1** | **G2A** | **Proteins** | **Acd** | **EtfB** | **EtfA** | **Hyp** |  |  |
|  |  |  | **Accession number** | WP_204662590 | WP_204662592 | WP_204662595 | WP_204662597 |  |  |

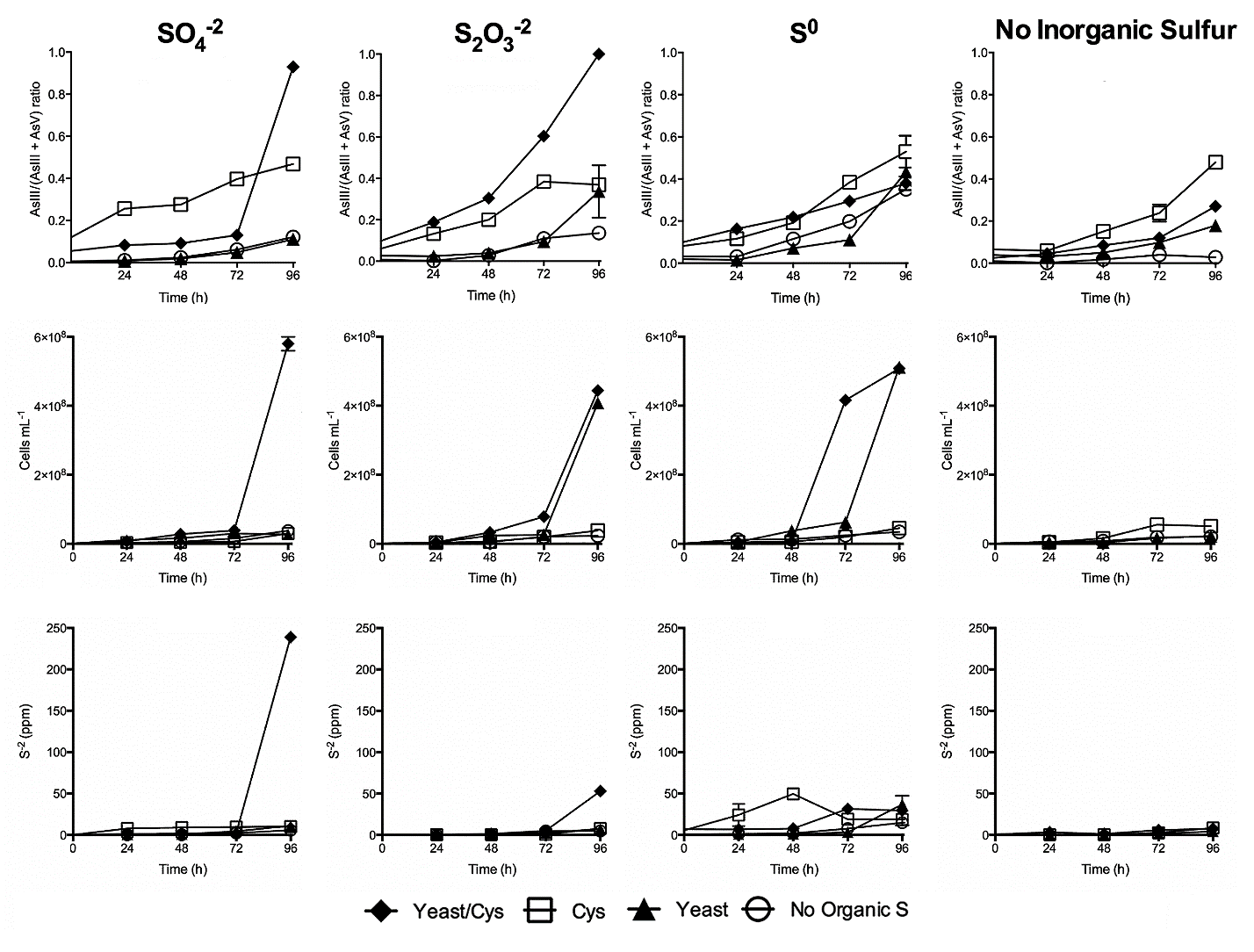

**Figure S1. *Fas* cultures amended with different inorganic and organic sulfur sources**. Sodium sulfate (SO_4_^-2^), sodium thiosulfate (S_2_O_3_^-2^), mineral sulfur (Sº) and no inorganic sulfur sources were tested, with yeast extract/cysteine (filled rhombus), cysteine (empty square), yeast extract (filled triangle), and no organic sulfur sources (empty circle). Graphs show arsenic ratio (AsIII/[AsIII + AsV]), cell number (cells mL^-1^) and S^-2^ production (ppm) in cultures performed in AsV (2 mM) Newman’s media with 1x10^6^ initial cells and incubated at 30 ºC in an anaerobic chamber under non-stirring conditions. Error bars represent the standard error of triplicate cultures.

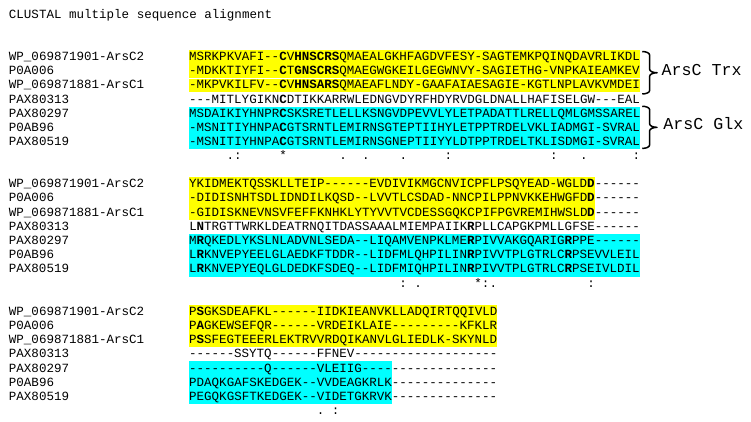

**Figure S2.** **Multiple alignment of aminoacid sequences of known and putative arsenate reductases dependent of glutaredoxins (Glx) or thioredoxin (Trx).** *E. coli* strain K12 (P0AB96), *Staphylococcus aureus* (P0A006), *Fas* (WP_069871881, ArsC-1 and WP_069871901, ArsC-2) and *Citrobacter* sp. TSA-1 (PAX80297; PAX80313 and PAX80519).

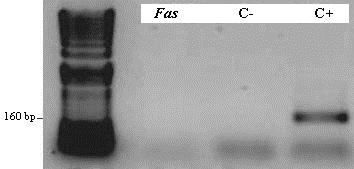

**Figure S3. Confirmation of the absence of the *arrAB* gene cluster in *Fas*.** PCR amplification using the primers arrAf and arrAr to target a ∼160–200 bp fragment of *arrA* gene was performed^[1]^. The negative control (C-) does not contain DNA template and, as positive control (C+), *Shewanella* sp. strain ANA-3 DNA was used.

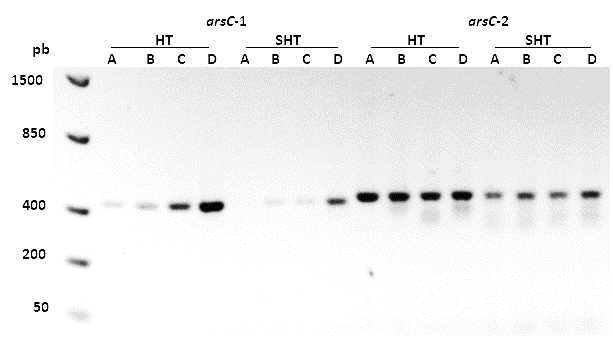

**Figure S4. PCR amplification of the arsenate reductase genes *arsC*-1 and *arsC*-2 from *Fas* genomic DNA at different hybridization temperatures**. (A) 60.0, (B) 59.2, (C) 58.0, and (D) 56.1 °C. The DNA sizes (bp) are indicated on the left side of the DNA ladder (first lane). SHT refers to products without His-tag, and HT to those with His-tag.

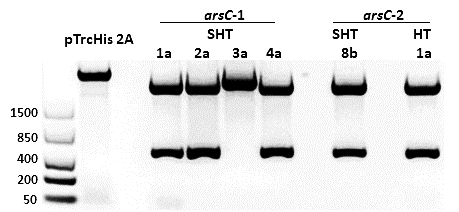

**Figure S5. Releasing of *arsC*-1 and *arsC*-2 DNA fragments from pGEM-T vector.** DNA fragments were released from different recombinant clones by digestion of DNA with *Xho*I and *Hind*III and purified from the agarose gel. The plasmid pTrcHis 2A was digested by the same restriction enzymes to allow the directional insertion of the *arsC* fragments. The DNA sizes (bp) are indicated in the left side of the DNA ladder (first lane).

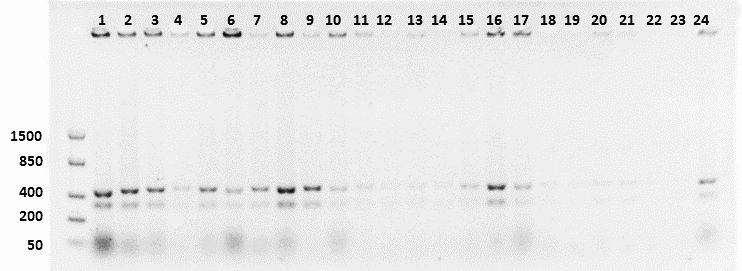

**Figure S6. *E. coli* WC3110 harboring the recombinant *arsC*-1*_Fas_* SHT expression vector checked by PCR.** The DNA sizes (bp) are indicated on the left side of the DNA ladder lane. Although not clearly seen in the photograph, all the 24 tested clones were positive.

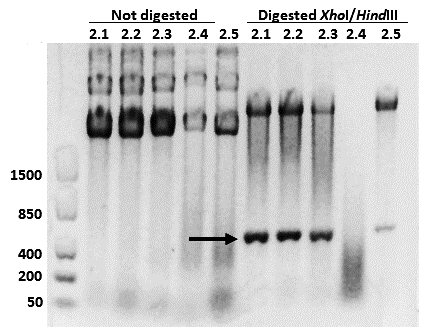

**Figure S7. Determination of the presence of the insert *arsC*-2*_Fas_* SHT by digestion of the plasmid pTrcHis2A.** The number on the head of each lane identifies the *E. coli* WC3110 recombinant clone from which the plasmid was obtained. The DNA sizes (bp) are indicated in the left side of the DNA ladder lane.

1. Malasarn, D. et al., arrA Is a Reliable Marker for As(V) Respiration. Science, 2004. 306(5695): p. 455.
